# Limited microglial metabolic improvement with time-restricted feeding in diet-induced obesity

**DOI:** 10.1101/2024.09.04.611250

**Authors:** Han Jiao, Jarne Jermei, Xian Liang, Hendrik J.P. van der Zande, Frank Vrieling, Valentina Sophia Rumanova, Milan Dorscheidt, Felipe Correa-da-Silva, Anhui Wang, Ewout Foppen, Bob Ignacio, Alberto Pereira Arias, Dirk Jan Stenvers, Tiemin Liu, Kimberly Bonger, Rinke Stienstra, Zhi Zhang, Andries Kalsbeek, Chun-Xia Yi

## Abstract

Time-restricted eating has shown promise for improving metabolic health in obese humans via incompletely resolved mechanisms. In this study, we investigated how time-restricted feeding (TRF) at different times of the day affects microglial immunometabolism using Wistar rats. We found that in high-fat diet (HFD)-fed obese rats, TRF during the active phase reduced fat mass, altered rhythmicity of the microglial transcriptome, and prevented an increase in hypothalamic microglia. These effects were dampened or absent with TRF during the resting phase. However, a HFD-induced microglial immunometabolic phenotype, characterized by reduced electron transport chain and increased lipid metabolism gene expression, and metabolic inflexibility, was not reversed by TRF in either the active or resting phase, indicating that reprogrammed microglial metabolism in obesity is a persistent cellular functional change that requires further study.

## Main

The growing global obesity crisis and its associated complications pose major health concerns [1–3]. Our modern society is associated with prolonged eating times and shorter fasting periods during the day-night cycle, which contributes to the development of obesity [4]. Time-restricted eating (TRE), by limiting the eating time window and increasing the fasting period, has emerged as a novel strategy for preventive healthcare and metabolic disease management [5–12]. Both human and animal studies have demonstrated that alignment of the time restricted eating (or feeding in animal studies) period with the body’s natural circadian rhythms is a crucial prerequisite for its beneficial effects [5, 8, 11, 13–17].

The immune activity of peripheral myeloid cells and brain microglia plays an important role during obesity development [18–24]. These cells tightly follow a circadian rhythmicity [25, 26], as *e.g.* the intrinsic core clock genes regulate key immune transcription factors that drive rhythmic expression of cytokines and immune receptors [26–31]. In addition, circadian proteins regulate inflammatory responses and immune cell trafficking [28, 29, 31–34]. Previous studies have shown that the rhythmicity of clock genes and immunometabolic pathways in microglia are disturbed when rats are subjected to an obesogenic high-fat diet (HFD) [35], possibly due to the fact that when fed a HFD their eating time is not restrained anymore to the active (dark) period [36], but also extends into the resting (light) period when these nocturnal animals usually sleep [5, 37].

The current study investigated whether time-restricted feeding (TRF) can reverse these obesity-associated microglial disturbances. Here, the impact of 10h-TRF on microglial function in male Wistar rats fed either a standard chow diet or HFD was assessed. Food access was restricted to either the dark phase (TRF dark-fed) or the light phase (TRF light-fed) and results were compared to animals without any time restriction, i.e. *ad libitum*-fed. After 4 weeks, TRF sacrifices were performed every four hours over the 24-hour light/dark cycle. Subsequently, brains and peripheral blood mononuclear cells (PBMCs) were isolated. Brains were used to isolate microglia for RNA sequencing, to profile microglia at the morphological level, and to measure cellular metabolism (Fig. 1a). Moreover, how TRF influences adiposity, the rhythmicity of microglial gene expression and their immune profiles were also assessed. Our results provide further insights into how feeding time alignment may alter HFD-induced disruptions in microglial immunometabolism and reveal potential pathways for managing diet-related metabolic disorders.

**Fig. 1.**
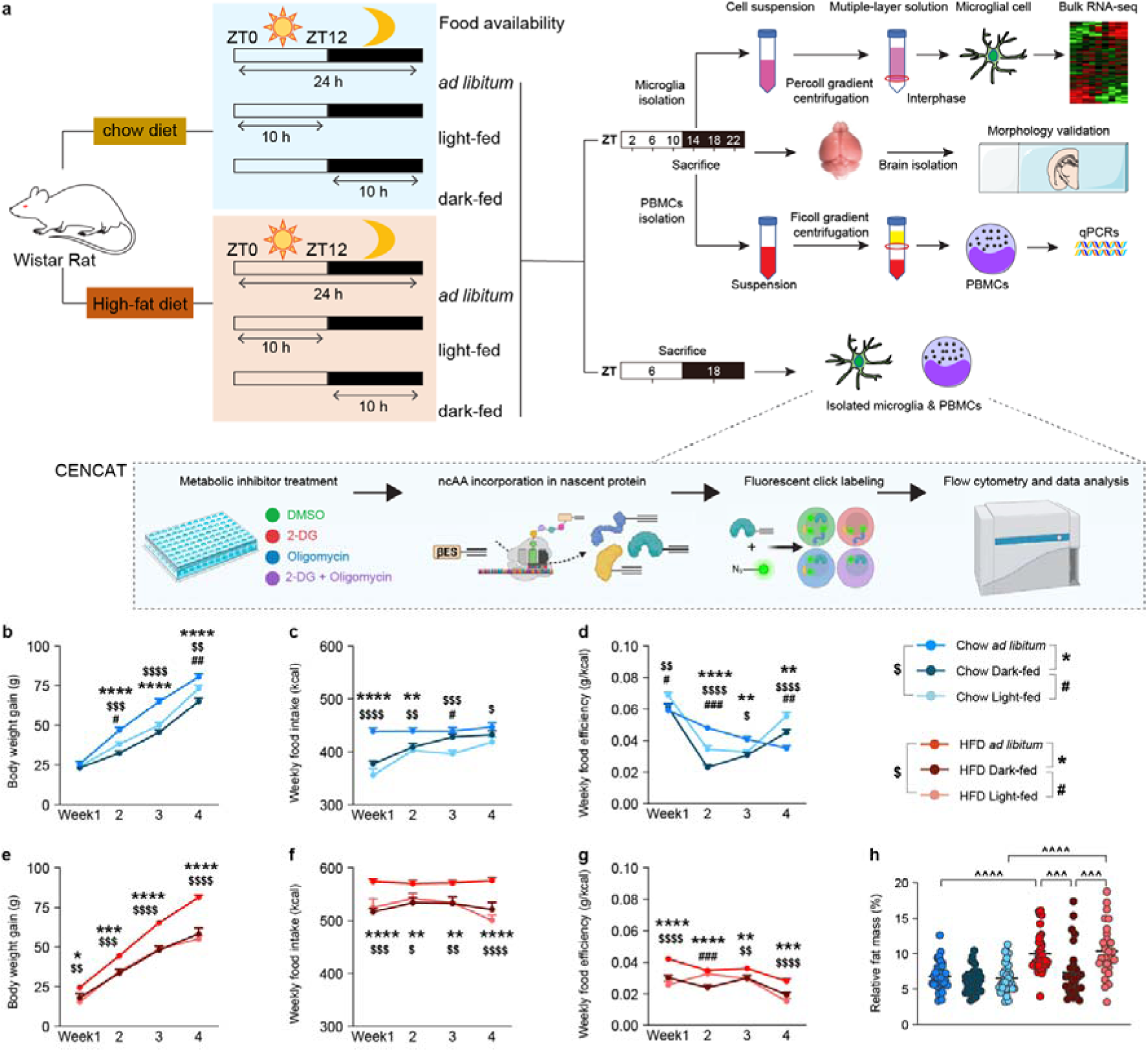
Time-restricted feeding with chow diet or high-fat diet differentially affects body weight and adiposity. **a**, Scheme illustrates the experimental design, showing the time-restricted access of different groups of Wistar rats to a normal chow diet or a high-fat diet (HFD) during 24 h, and the strategies used to isolate microglia and peripheral blood mononuclear cells (PBMCs) and for bioinformatic and morphological analyses. After sample processing and plating, functional metabolic profiles were determined using CENCAT (Cellular Energetics through Non-Canonical Amino acid Tagging). ZT0D=Dlights on, ZT12D=Dlights off; Light-fed: ZT1-ZT11 (10h), Dark-fed: ZT13-ZT23 (10h), *ad libitum*: free access to food (24h). β-ethynylserine (βES). **b-g**, The dynamic changes in weekly body weight gain, food intake, and food efficiency under *ad libitum*, light-fed, and dark-fed conditions on chow diet (**b, d, f**) or high-fat diet (HFD) (**c, e, g**). **h**, Relative fat mass in all conditions just before sacrifice. Data are shown as mean ± SEM, data and were analysed by two-way ANOVA with repeated measures (*, #,

## Results

### TRF decreases body weight gain and differentially modulates adiposity

Given that rats are nocturnal animals, feeding them during the dark phase aligns with their natural active period[5, 8]. We hypothesize that time-restricted feeding (TRF) in the dark phase may enhance circadian rhythms and metabolic health, potentially leading to more favourable body parameter outcomes and circadian rhythm compared to feeding during the light phase, which coincides with their resting period. Based on this hypothesis, we first explored the effects of TRF during both the light and dark phases on body weight, food intake, and adiposity. We found that in the standard chow diet condition, both TRF light- and TRF dark-feeding reduced body weight gain in the second and third week compared to the *ad libitum* group. However, this impact diminished in the fourth week (Fig. 1b). This is likely explained by the *ad libitum*-fed rats consistently showing higher caloric intake during the first 3 weeks of the TRF period (Fig. 1c). Moreover, food efficiency decreased over time in the *ad libitum* fed animals but was higher than both TRF groups in week 2 and 3 (Fig. 1d). In contrast, after an initial drop in week 2, food efficiency increased in both TRF groups, with the TRF light-fed group displaying a significantly higher food efficiency in week 4 (Fig. 1d). This indicates increased body weight gain with the same amount of food intake in TRF light-fed rats compared to TRF dark-fed and *ad libitum* fed rats.

In the HFD condition, both TRF groups showed lower body weight gain compared to the *ad libitum* group (Fig. 1e). Similar to the chow-fed condition, HFD *ad libitum* rats exhibited increased caloric intake (Fig. 1f), but in contrast both the TRF light- and dark-fed group exhibited lower food intake and food efficiency compared to the *ad libitum* group during the entire experiment (Fig. 1f, 1g). As expected, HFD significantly increased relative fat mass in Wistar rats (Fig. 1h). No difference in adiposity was found between the three chow-fed groups (Fig. 1h). However, in the HFD condition, adiposity was reduced in TRF dark-fed rats compared to both *ad libitum* and TRF light-fed rats (Fig. 1h), as also described by others [5]. This suggests that even with an unhealthy diet, aligning the daily feeding period with the active phase can reduce adiposity.

### HFD restricted to the dark phase significantly reduces microglial cell number in the ARC

The arcuate nucleus (ARC) of the hypothalamus plays a critical role in regulating systemic energy homeostasis [38]. It is known that a high-fat diet significantly increases the number and ramifications of microglia in the hypothalamus compared with a chow diet [39, 40]. Thus, we performed immunohistochemistry targeting ionized calcium-binding adaptor molecule 1 (Iba1) as a microglial marker to assess cell numbers and ramifications (Fig. 2a). As expected, the obesogenic diet caused an increased microglial cell number and ramifications in the hypothalamus in HFD *ad libitum* and HFD light-fed group as compared to the chow *ad libitum* condition (Fig. 2b,2c). Remarkably, we found that this increase in the number of microglial cells was not observed in the HFD dark-fed group (Fig. 2b), whereas TRF did not affect the number of ramifications. We also investigated the spatial relationship between neurons and microglia across the circadian cycle (Fig. 2d-2f). In the chow diet group, microglia were found to be in closer proximity to neurons during the night (ZT18) compared to the day (ZT6), indicating a circadian modulation of neuron-microglia interactions (Fig. 2e,2f). However, in the HFD group, this circadian variation in microglial-neuronal contact was absent, with microglia consistently maintaining close proximity to neurons, irrespective of the time of day (Fig. 2e,2f). TRF did not significantly alter neuron-microglia proximity in neither chow nor HFD-fed rats (Fig.2e,2f).

**Fig. 2.**
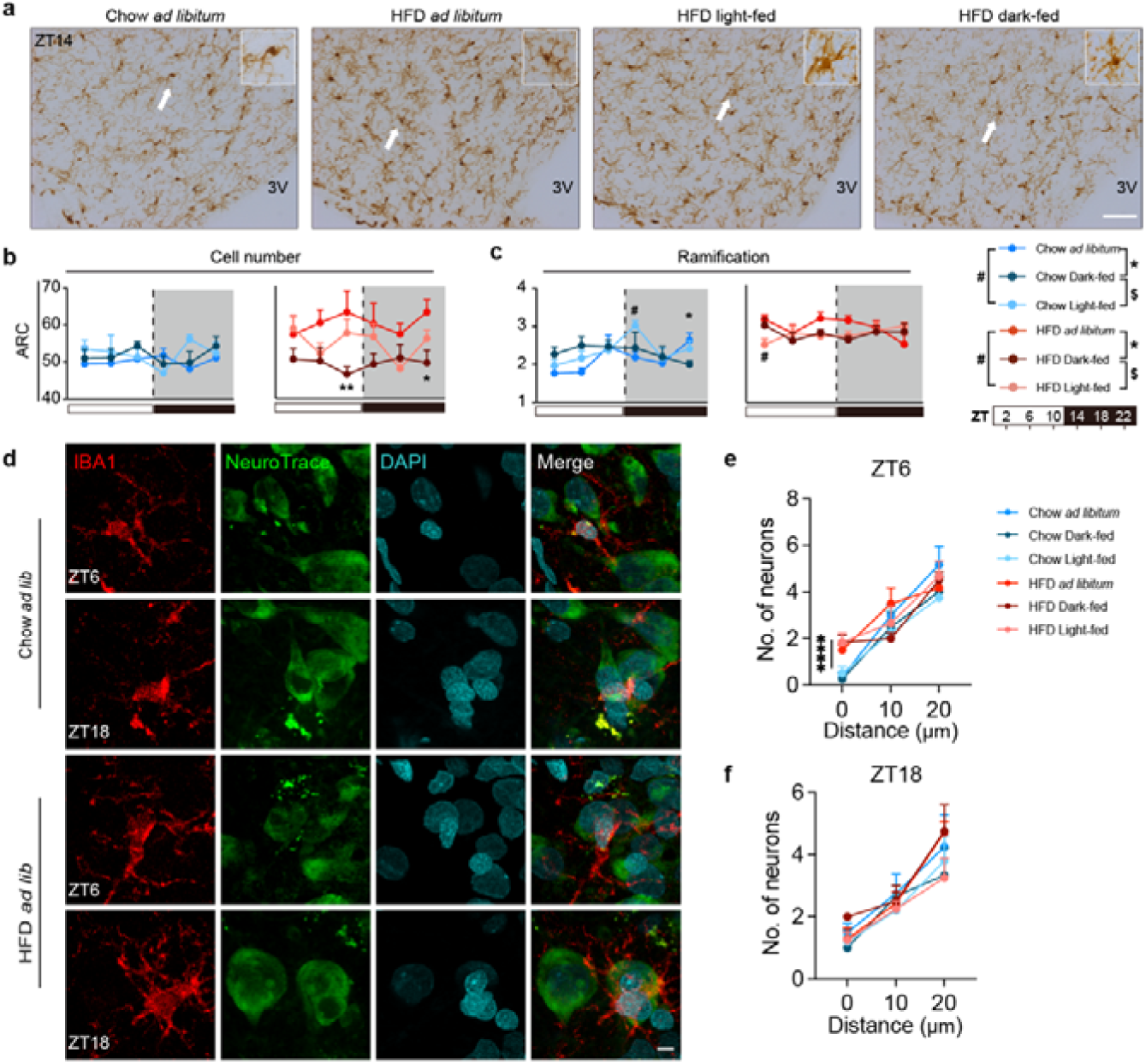
HFD restricted to the dark phase significantly reduces microglial cell number in the ARC. **a**, Representative figures of Iba1-ir microglia in the arcuate nucleus (ARC) at ZT14. The microglial cells in the high-magnification images in the upper right corner are indicated by a white arrow in the overview image. **b,c,** Quantifications of microglial cell number (b) and ramifications (c) in the ARC (0.16 mm^2^). Scale bar = 50 μm. 3V = the third ventricle. Iba1= Ionized calcium binding adaptor molecule 1. **d**, Representative confocal images showing colocalization of Iba1 (red, microglial marker), NeuroTrace (green, neuronal marker) and DAPI (cyan, nuclear marker) in the hypothalamus across different dietary conditions (Chow *ad libitum*, HFD *ad libitum*) at circadian time points ZT6 and ZT18. Scale bar: 5 µm. **e,f,** Quantification of the spatial distance between neurons and microglia in the chow diet and HFD diet groups. Data are shown as mean ± SEM, P values are analysed by two-way ANOVA with repeated measures. Statistical significance is indicated as *p < 0.05, **p < 0.01.

### Microglial morphological characteristics in other brain regions

We also wondered whether HFD and TRF affected microglial morphological characteristics across distinct brain regions, including the cortex, hippocampal CA1, thalamus and ventromedial nucleus of the hypothalamus (VMH), or whether these alterations are ARC-specific (Extended Data Fig. 1). Interestingly, we observed an increase in the number of microglia in the HFD dark-fed group compared to the HFD *ad libitum* group at the beginning of the light phase in the cortex, hippocampal CA1, and thalamus regions. In addition, light-fed TRF reduced the number of VMH microglia at the onset of the light period compared to the *ad libitum* group in HFD-fed rats. This observation implies that the timing of feeding may affect microglial cell numbers differentially across various brain areas. We also found that HFD increased the number of microglial ramifications across all brain areas investigated, which was not prevented by changes in (TRF-induced) feeding patterns. Together these data indicate that the reduced microglial cell numbers in HFD dark-fed rats is ARC-specific, and that microglial ramifications (as a potential proxy for microglial activity or inflammation) increase with HFD irrespective of TRF.

### TRF alters oscillatory expression of microglial core clock genes

To assess whether TRF functionally alters microglia, we first examined the impact of TRF on the oscillatory expression of molecular clock genes in microglia. We found that most clock gene rhythms were preserved in both chow light-fed and dark-fed groups compared to the chow *ad libitum* group, but rhythmicity in *Cry1* and *Per1* was lost in the chow light-fed group (Fig. 3a, 3c). Additionally, TRF in the light-fed group significantly altered the acrophase of *Arntl*, *Nr1d1*, *Clock*, and *Rora* compared to the chow ad libitum group (Fig. 3b, 3d), while TRF in the dark-fed group strengthened the amplitude of *Rora* expression relative to the chow *ad libitum* group (Fig. 3a, 3b). In the HFD condition, all clock genes showed significant daily oscillations in both the *ad libitum* and TRF dark-fed groups (Fig. 3e, 3g), while the TRF light-feeding strategy caused a loss of rhythmicity in *Clock*, *Rora* and *Cry2* (Fig. 3e, 3g). The amplitude of *Cry2* oscillation were higher in the HFD dark-fed group compared HFD *ad libitum* (Fig. 3g,3h). In summary, TRF dark-feeding significantly strengthened the amplitude of certain clock genes, while in the TRF light-fed group, rhythmic expression was maintained in general, but the amplitude and acrophase of most clock gene oscillations were significantly altered compared to *ad libitum,* both in the chow and HFD conditions (Fig. 3).

**Fig. 3.**
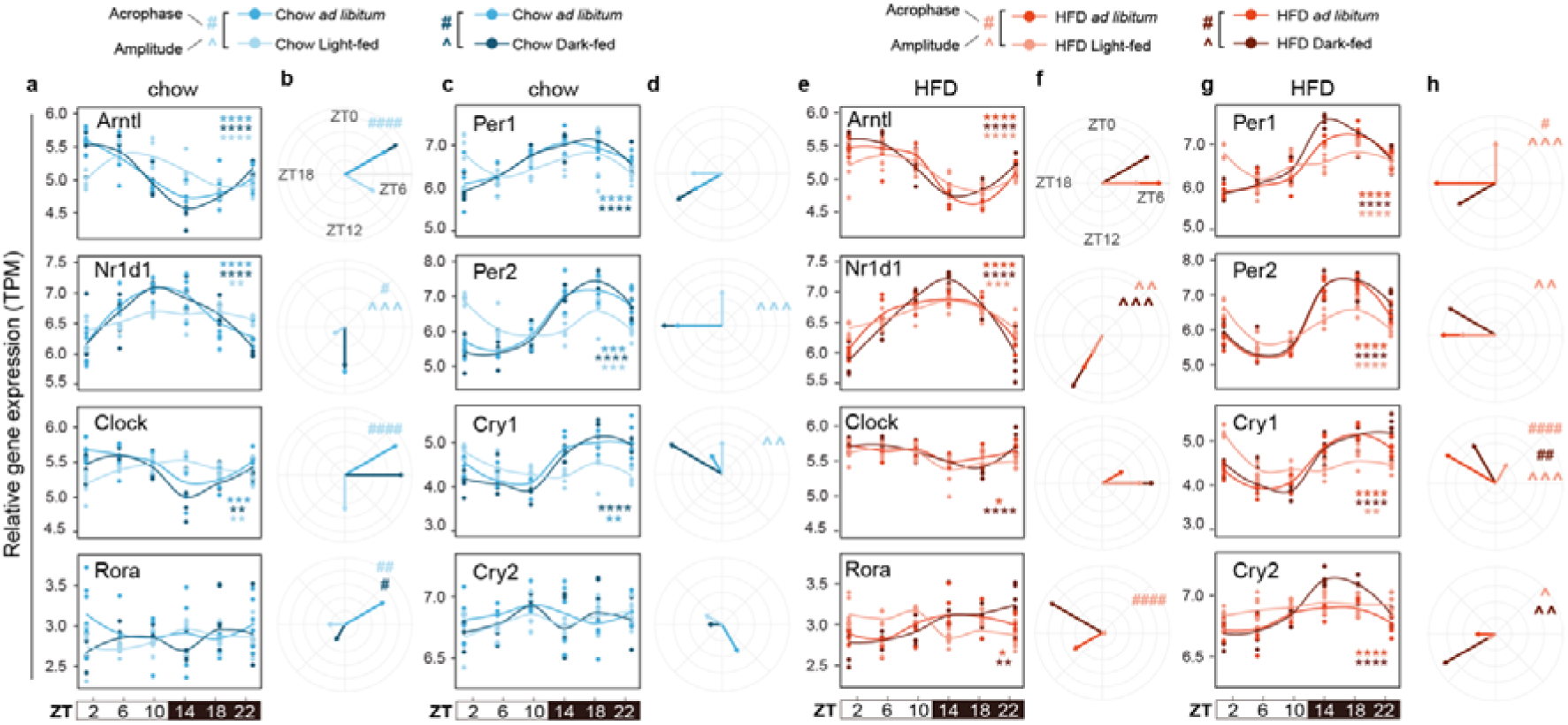
TRF during the light phase alters the acrophase and/or amplitude of microglial core clock gene expression. a,c,e,g, Relative gene expression of core clock genes in different feeding patterns under Chow **(a,c)** and HFD **(e,g)** conditions over time (timepoints from ZT2 to ZT22, at 4-hour intervals). Coloured asterisks in the upper or lower right corner indicate a significant 24-hour rhythm according to JTK_CYCLE. *P < 0.05, **P < 0.01, ***P < 0.001, ****P < 0.0001. **b,d,f,h,** Spider plots show acrophase (coloured arrow) and amplitude (length of the coloured arrow) alterations of core clock genes under Chow **(b,d)** and HFD **(f,h)** conditions. Coloured carets (^) and number signs (#) represent significant differences in amplitude and acrophase, respectively. ^#, ^^P < 0.05, ^##, ^^^P < 0.01, ^###, ^^^^P < 0.001, ^####^P < 0.0001.

In obesity, circulating immune cells are recruited to peripheral metabolic tissues such as adipose tissue, contributing to inflammation and insulin resistance [41–43]. Given our observations in microglia, we thus wondered whether peripheral blood mononuclear cells (PBMCs) also displayed altered molecular clock gene oscillations upon TRF. However, core clock gene oscillations were largely unaltered in total PBMCs, suggesting this is a microglia-specific response to TRF (Extended Data Fig. 2).

### Microglial gene rhythmicity is highly sensitive to TRF in chow-fed rats

To compare alterations in microglial transcriptional rhythmicity in rats under different feeding conditions we employed the CompareRhythm R package [44], which uses probability assignment to categorize rhythmicity alternations into ‘gain’, ‘loss’, or ‘changed’ rhythmicity, as well as identifying genes with ‘same’ rhythmicity within a 24-hour timeframe (Fig. 4). Evaluation of the number of rhythmic genes in the different groups showed the following: 2,142 rhythmic genes in the chow *ad libitum*-fed group, 4,109 in the chow dark-fed group, 3,910 in the chow light-fed group, and 3,805 in the HFD *ad libitum*-fed group, 4,623 in the HFD dark-fed group, and 1,770 in the HFD light-fed group (Fig. 4a). This suggests that in HFD-fed rats, light-restricted TRF markedly decreases gene rhythmicity in microglia as compared to chow-fed rats.

**Fig. 4.**
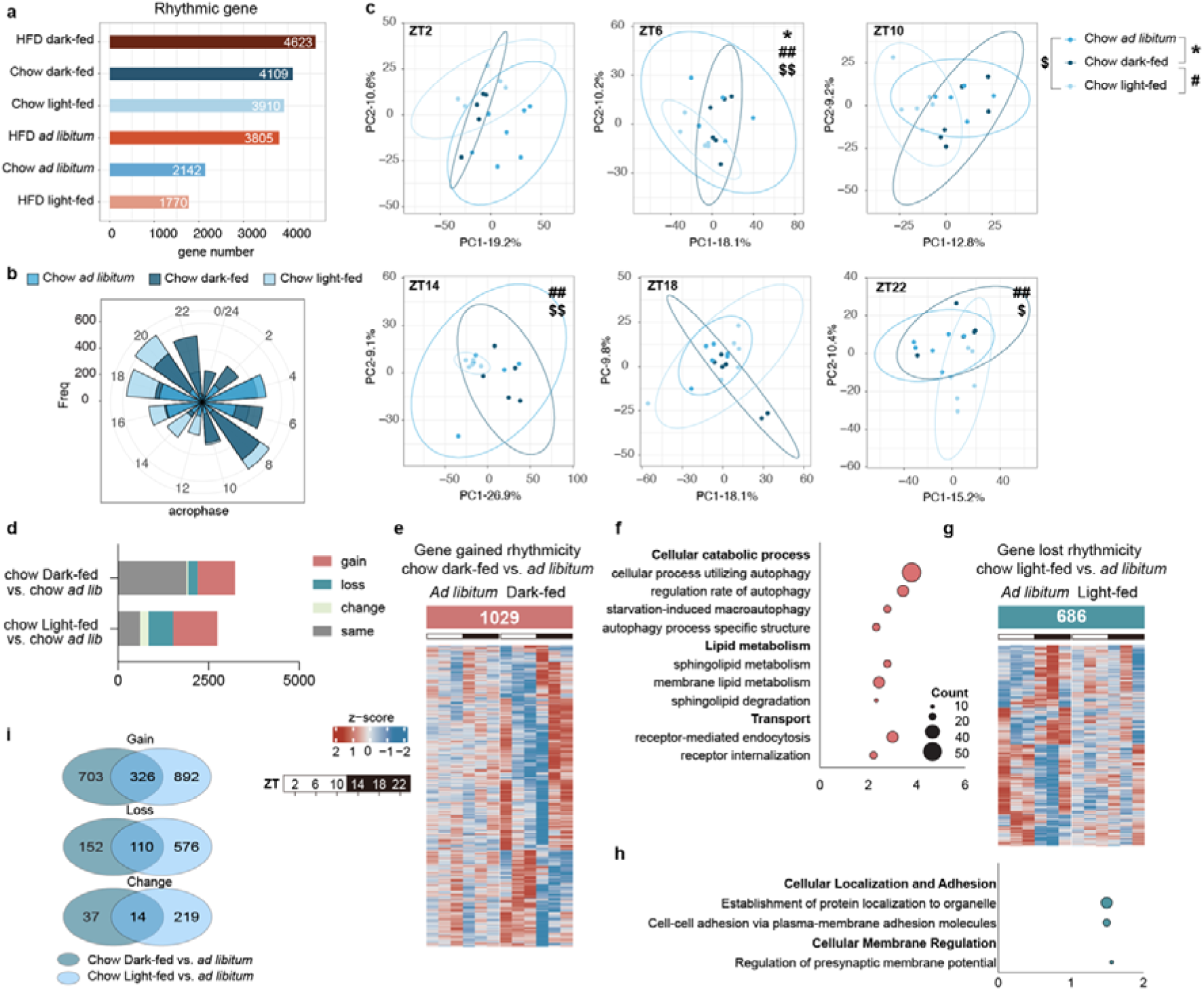
Light- and dark-phase TRF uniquely alters gene expression rhythms in rats fed a normal chow diet. a, The number of rhythmic genes in each group. **b**, Acrophase plot of all transcripts after rhythmicity analysis (bars represent the number of genes exhibiting their acrophase at each time point). **c,** Principal component analysis (PCA) of the microglial transcriptome in each of the time points assessed. Data analyzed using PERMANOVA. **d,** (Stacked) bar graph depict changed rhythmicity categories of gained, lost, changed, same and arrhythmic genes in different comparisons in the chow-fed groups. Ge ne numbers are shown on the x-axis. **e**, Heatmap of gained rhythmicity genes in chow dark-fed vs chow *ad lib* group. Unfilled rectangles showed the number of gained rhythmicity genes. Heatmaps display the z-scored normalized transcript expression levels in different comparisons. Transcript expression profiles were categorized based on differential rhythmicity analysis using CompareRhythmR [44]. **f**, Pathways involved gained rhythmicity genes in the chow dark-fed vs. chow *ad libitum* group. Enlarged and bold terms serve as summaries of sub pathways. Dot size corresponds to the number of genes associated with each pathway, and log10 adjusted p-value thresholds are shown on the x-axis. **g**, Heatmap of lost rhythmicity genes in chow light-fed vs chow *ad lib* group. **h**, Pathways involved lost rhythmicity genes in the chow light-fed vs. chow *ad libitum* group. **i**, Venn diagrams illustrate the overlapping number of genes with gained, lost, and changed rhythmicity in chow comparisons.

In chow diet-fed rats, both light-fed and dark-fed groups displayed distinct rhythmic transcriptomic patterns compared to chow *ad libitum*, with notable shifts in acrophase distribution (Fig. 4b). Principal component analysis (PCA) combined with PERMANOVA revealed significant effects on the microglial transcriptome of light-feeding at ZT6, ZT14, and ZT22, and of dark-feeding at ZT6 (Fig. 4c). Light-feeding was particularly disruptive to microglial gene expression, with approximately 76% of rhythmic genes showing gain, loss, or a change in rhythmicity (Fig. 4d). In contrast, dark-feeding maintained rhythmicity in around 60% of genes compared to the *ad libitum* group (Fig. 4d).

Further analysis showed that genes which gained rhythmicity in the chow dark-fed group, which is fed during the active phase, compared to chow *ad libitum* (Fig. 4e) were functionally enriched in processes such as cellular catabolism, lipid metabolism, and transport (Fig. 4f). Conversely, genes that lost rhythmicity in the chow light-fed group, which is fed during the resting phase, compared to chow *ad libitum* (Fig. 4g) were enriched in general cellular processes related to cellular localization, adhesion, and membrane regulation (Fig. 4h). In the chow condition, light-fed and dark-fed rats shared 326 genes with lost rhythmicity, whereas 110 genes that gained rhythmicity were shared, and 14 that shifted rhythmicity compared to HFD *ad libitum* (Fig. 4i). Overall, in chow-fed rats, both TRF regimens induced clear adaptive changes in microglial gene expression.

### TRF light-fed group displayed significant loss of microglial gene rhythmicity in HFD-fed rats

During HFD-feeding, the dark-fed group displayed increased rhythmic microglial transcriptomic patterns compared to the HFD *ad libitum* group, with notable increased frequencies in acrophase distribution at mid-day and mid-night (Fig. 5a, 5c). On the contrary, fewer genes were rhythmic in microglia from the HFD light-fed group (Fig. 4a), and their acrophase distribution additionally shifted towards one peak during the light phase (Fig. 5a). PCA analysis combined with PERMANOVA revealed significant effects on the microglial transcriptome of both light-feeding and dark-feeding only at ZT2 (Fig. 5b). Light-feeding was particularly disruptive to microglial gene expression, as nearly 2000 genes lost their rhythmic expression in the TRF light-fed group compared to the *ad libitum* group (Fig. 5c). In contrast, dark-feeding gained rhythmicity in 775 genes compared to the *ad libitum* group (Fig. 5c), while the majority of rhythmic genes were unchanged.

**Fig. 5.**
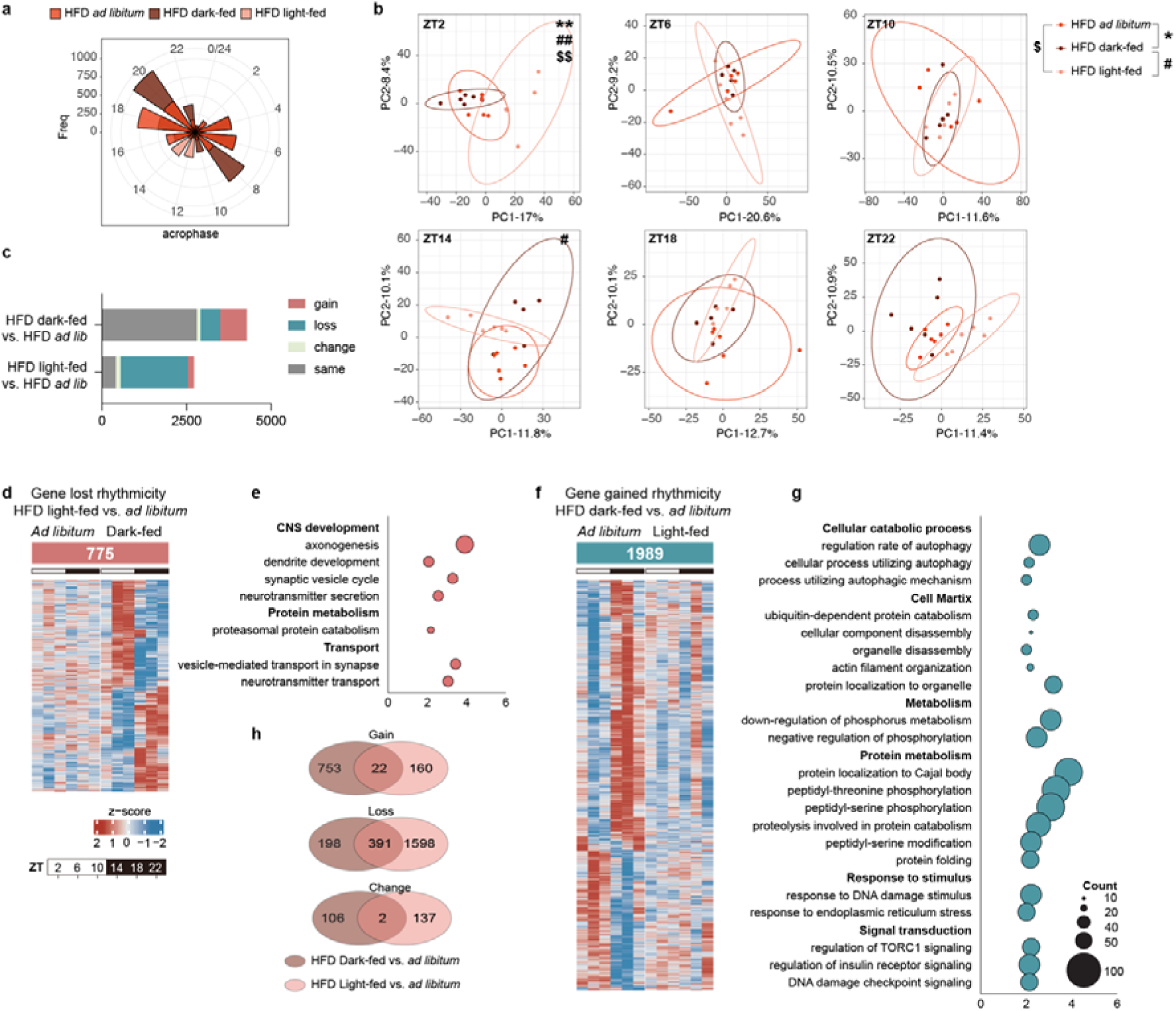
Time-restricted feeding in the light-phase combined with HFD significantly reduces rhythmicity in microglial gene expressions. **a**, Acrophase plot of all transcripts after rhythmicity analysis (bars represent the number of genes exhibiting their acrophase at each time point). **b,** Principal component analysis (PCA) of the microglial transcriptome in each of the time points assessed. Data analyzed using PERMANOVA. **c,** (Stacked) bar graph depict changed rhythmicity categories of gained, lost, changed, same and arrhythmic genes in different comparisons in the HFD-fed groups. **d**, Heatmap of gained rhythmicity genes in HFD dark-fed vs HFD *ad lib* group. **e**, Pathways involved gained rhythmicity genes in the HFD dark-fed vs. HFD *ad libitum* group. **f**, Heatmap of lost rhythmicity genes in HFD light-fed vs HFD *ad lib* group. **g**, Pathways involved lost rhythmicity genes in the HFD light-fed vs. HFD *ad libitum* group. **h**, Venn diagrams illustrate the overlapping number of genes with gained, lost, and changed rhythmicity in HFD comparisons.

Further analysis showed that genes which gained rhythmicity in the HFD dark-fed group, which is fed during the active phase, compared to HFD *ad libitum* (Fig. 5d) were functionally enriched in general cellular processes such as CNS development, protein metabolism and transport (Fig. 5e). Conversely, genes that lost rhythmicity in the HFD light-fed group, which is fed during the resting phase, compared to HFD *ad libitum* (Fig. 5f) were enriched in cellular catabolic process, cell matrix, metabolism, protein metabolism, response to stimulus and signal transduction (Fig. 5g). In the HFD condition, light-fed and dark-fed rats shared 391 genes with lost rhythmicity, whereas only 22 genes that gained rhythmicity were shared, and 2 that shifted rhythmicity compared to HFD *ad libitum* (Fig. 5h).

### HFD significantly impairs the elasticity of microglial pathways

We then investigated how HFD feeding affects the rhythmicity of microglia, as well as the potential for TRFs to mitigate these effects. We found that HFD significantly altered the rhythmicity of over 1200 genes compared to chow-fed controls, indicating that a high-fat diet reshapes the time-of-day-dependent expression of microglial genes (Extended Data Fig. 3a). Of note, we found approximately 55% of oscillatory genes showed no changes, whereas the rest mostly showed loss or gain of rhythmicity (Extended Data Fig. 3a,3b). The enriched pathway analyses show that genes with lost rhythmicity were mainly involved in cellular organization, CNS development, transport and immune response (Extended Data Fig. 3c), while genes with gained rhythmicity were mainly involved in protein metabolism (Extended Data Fig. 3d). Subsequently, we examined whether these HFD-induced alternations are affected by the HFD TRF dark-fed condition, which would suggest these HFD-induced disruptions are rescued by dark-fed TRF (Extended Data Fig. 3f). We found that a group of genes, marked in light blue in the Sankey diagram (Extended Data Fig. 3g), gained rhythmicity in HFD *ad libitum* vs chow *ad libitum*, whereas they lost their rhythmicity in the HFD dark-fed condition (Extended Data Fig. 3g,3h). Pathway enrichment analysis of these light blue-marked genes mainly revealed cell cycle-related pathways (indicated in red) and lipid metabolism pathways (indicated in blue) (Extended Data Fig. 3h), suggesting that TRF during the dark phase may mitigate HFD-induced disruption in microglial cell cycle and lipid metabolism rhythmicity.

Through unsupervised gene set variation analysis (GSVA) combined with the Mfuzz time clustering method [45, 46], we generated clusters of gene sets of which the expression followed similar patterns over time, to estimate the rhythmicity of pathway activity across the sample population, which yielded four distinct clusters (Extended Data Fig.4a). Notably, the chow dark-fed group constituted the largest proportion of pathways in cluster 4 (with a peak at the onset of the dark period), comprising nearly 80% of the samples (gene lists available in deposited data). We then studied variation in clusters among the three different feeding patterns, and found there was a marked transition of genes from cluster 3 in the chow *ad libitum* group to cluster 4 in the chow dark-fed group (Extended Data Fig.4c). Few shifts were observed from clusters 1-2 in the chow *ad libitum* group to clusters 1-3 in the chow light-fed group (Extended Data Fig.4b). The same analysis on HFD conditions revealed reduced pathway enrichment compared with chow conditions (Extended Data Fig. 4d). While no cluster shifting was observed between HFD *ad libitum* and light-fed groups (Extended Data Fig.4e), some transitions were still evident between HFD *ad libitum* and HFD *dark-fed* (Extended Data Fig.4f), for instance from HFD *ad libitum* cluster 1 to cluster 4 in HFD dark-fed, and from cluster 2 in HFD *ad libitum* to cluster 1 in HFD dark-fed (Extended Data Fig.4f). These results imply that genes display higher rhythmicity in chow-fed rats, and additionally that elasticity of gene rhythms is perturbed in HFD-fed rats.

### High-fat diet feeding causes an immunosuppressive or hyporesponsive state in microglia at ZT18, which was not prevented by TRF

Principal component analysis (PCA) further demonstrated that the overall microglial transcriptome is significantly altered in HFD-fed rats compared to chow-fed control at both ZT2 and ZT18 (Fig. 6a). While TRF reduces body weight gain in HFD-fed rats, PCA analysis reveals that it does not fully restore the microglial transcriptome to that of chow-fed controls, particularly at ZT18, indicating persistent alterations under HFD even with TRF intervention (Fig. 6a).

**Fig. 6.**
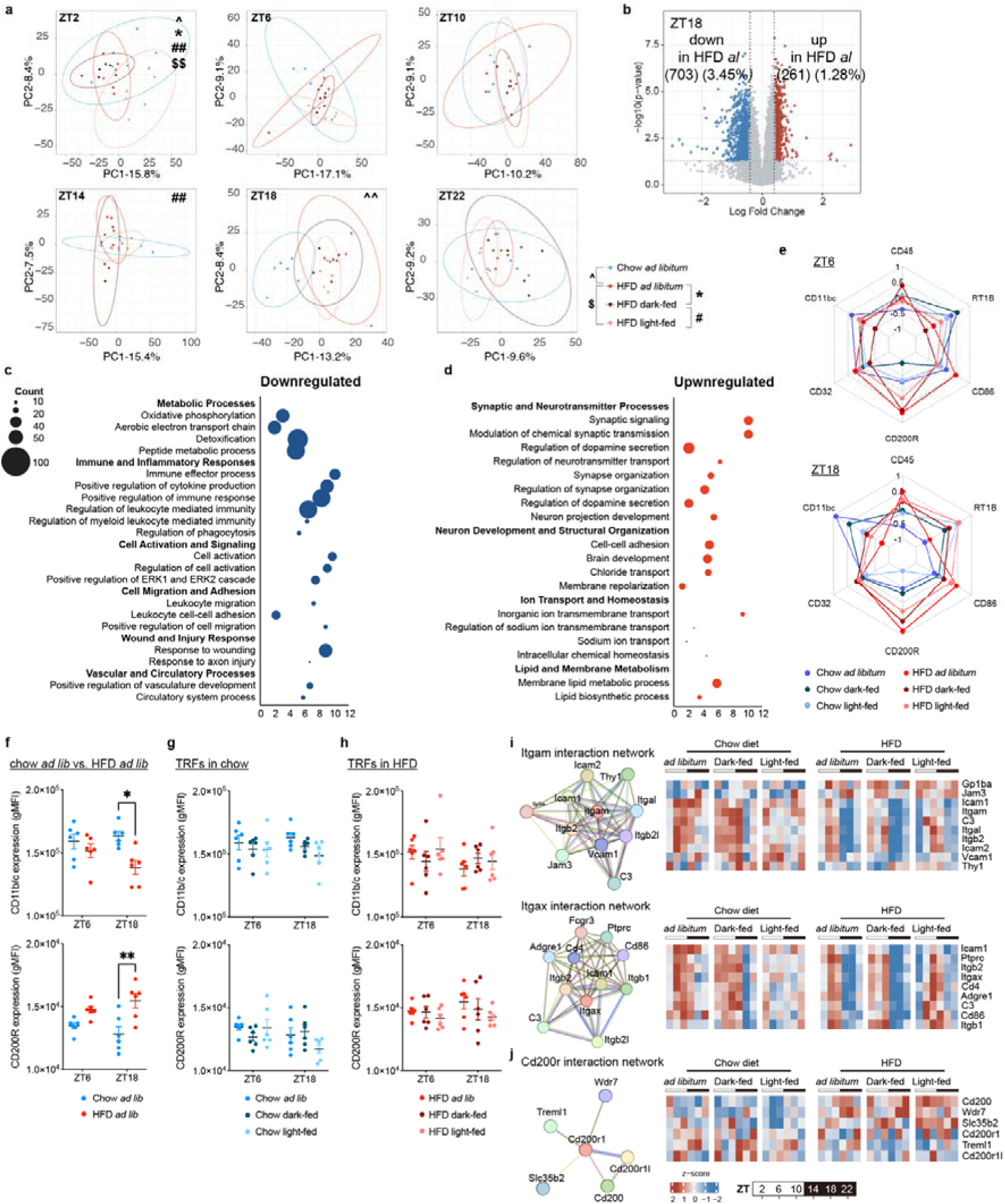
A high-fat diet-induced microglial immunophenotype is not reversed by TRFs. **a**, Principal component analysis based on variance-stabilized counts of all genes in chow *ad libitum*, HFD *ad libitum*, HFD dark-fed and HFD-light fed groups at all ZTs. Data analyzed using PERMANOVA. **b,** Volcano plots showing differentially expressed genes (DEGs) at ZT18 under chow *ad libitum* and HFD *ad libitum* conditions. Genes with significant changes (pD<D0.05) are indicated in red (fold change ≥D2) or blue (fold changeD≤D-2). **c,** Pathway enrichment analysis of up-regulated genes in HFD *ad libitum* compared to chow *ad libitum* at ZT18 using Metascape and showing enrichment in functions associated with synaptic signalling pathways. **d,** Pathway enrichment analysis of down-regulated genes in HFD *ad libitum* compared to chow *ad libitum* at ZT18 showing enrichment in functions associated with immune and inflammatory response. **e**, Spider plots depicting z-scored expression (geometric mean fluorescence intensity, gMFI) of indicated cell surface receptors on microglia at ZT6 (left) and ZT18 (right), determined via flow cytometry. **f-h**, gMFI of CD11b/c and CD200R in chow *ad libitum* vs. HFD *ad libitum* (**f**), different TRFs in chow diet (**g**) and different TRFs in HFD (**h**) at ZT6 and ZT18. Data are shown as mean ± SEM, P values are analysed by two-way ANOVA with repeated measures (* p< 0.05, **p<0.01). **i,j**, The protein-protein interaction network and genes associated with the network are given for Itgam, Itgad (**i**) and Cd200r (**j**), respectively.

We thus next investigated the impact of HFD on differential gene expression in microglia, and notably found a substantial downregulation of DEGs at ZT18 in the HFD group (Fig. 6b, Extended Data Fig.3e). The pathway analysis of downregulated genes showed enrichment mainly in metabolic processes (i.e. electron transport chain, ETC), immune & inflammatory response and cell activation (Fig.6c) and upregulated genes showed enrichment mostly in regulation of synaptic & neuronal process and lipid metabolism (Fig.6d). Flow cytometry confirmed these findings (Fig. 6e, and Extended Data Fig. 5a for microglial gating strategy), showing decreased surface expression of CD11b/c (a microglial activation marker) and increased CD200R (an inhibitory receptor) expression at ZT18 in the HFD group (Fig. 6f). Together, these data indicate that HFD induces an immunosuppressive or hyporesponsive state in microglia during the active phase. When analysing the effects of TRF in both chow and HFD groups, we observed no significant changes in microglial activation markers (Fig. 6g,6h), suggesting that TRF may have limited ability to reverse the microglial immunophenotype in HFD-fed rats. Additionally, protein-protein interaction analysis of CD11b/c and CD200R networks showed downregulation of genes associated with CD11b/c and upregulation of those linked to CD200R under HFD conditions, further supporting the HFD-induced shift in the microglial immune phenotype, which is unaffected by TRF (Fig. 6i,6j).

### Differential effects of TRF on microglial immunometabolism upon chow and HFD feeding

Immune cell function is regulated by their cellular metabolism. The microenvironment of immune cells may therefore impact their effector functions, through dynamic reprogramming of intracellular metabolic pathways [47]. Microglial function is also known to be highly dependent on metabolic flexibility, allowing them to shift between resting and active states by leveraging energy synthesis pathways like glycolysis, oxidative phosphorylation (OXPHOS), lipid metabolism, and amino acid metabolism [33, 46]. Our RNA sequencing analysis revealed significant downregulation of electron transport chain and lipolysis genes, and upregulation of lipid uptake and lipogenesis genes in HFD-fed rats compared to chow-fed rats (Fig. 7a, 7b). While TRF altered rhythmicity of lipid uptake and lipogenesis genes, expression levels were still markedly increased compared to chow-fed rats, suggesting significant HFD-induced disruption of mitochondrial metabolism that is not restored by TRF.

**Fig. 7.**
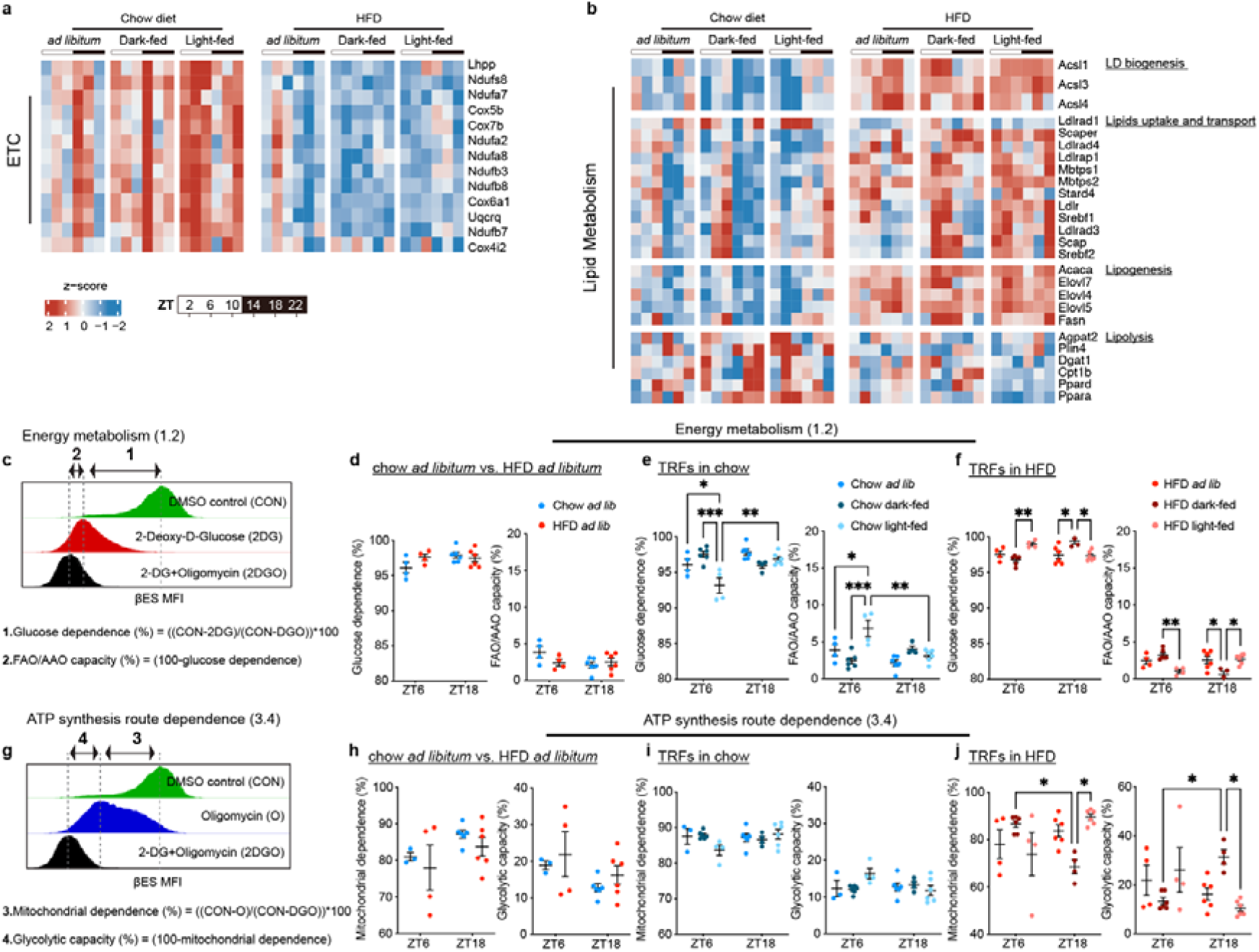
TRFs distinctly alter **microglial metabolism dependent on the diet fed. a,b,** Heatmaps depict z-scored normalized transcript expression levels of genes related to the electron transport chain (ETC), and lipid metabolism. LD, lipid droplets. **c,** Illustration of energy metabolism from glucose dependence (1) or FAO/AAO capacity (2) by calculation according to the following formulas [86]. Mean fluorescence intensities (MFI). **d-f**, Energy metabolism of microglia using CENCAT from chow *ad libitum* vs. HFD *ad libitum*-fed rats (**d**), different TRFs in chow diet-fed rats (**e**) and different TRFs in HFD-fed rats (**f**) at ZT6 and ZT18. **g,** Illustration of ATP produced from mitochondrial dependence (3) or glycolytic capacity (4) by calculation according to the following formulas [86]. **h-j**, Dependence on route of ATP synthesis of microglia from chow *ad libitum* vs. HFD *ad libitum-*fed rats (**h**), different TRFs in chow diet-fed rats (**i**) and different TRFs in HFD-fed rats (**j**) at ZT6 and ZT18. Data are shown as mean ± SEM, P values are analysed by two-way ANOVA with repeated measures (* p< 0.05, **p<0.01, ***p<0.01).

Seahorse XF analysis has been crucial in immunometabolism research by measuring cellular oxygen consumption (OCR) and extracellular acidification (ECAR) to assess metabolic profiles. However, it requires purified cells, high cell counts, and specialized equipment and reagents. To study in more detail the impact of HFD and TRF on microglial metabolism, we employed CENCAT (Cellular Energetics through NonCanonical Amino acid Tagging) [48]. CENCAT uses fluorescent click-labelling of the alkyne-bearing non-canonical amino acid β-ethynylserine (βES) to monitor protein synthesis via flow cytometry, serving as a proxy for metabolic activity. Assessing protein synthesis under conditions of metabolic inhibition, *i.e.* inhibiting glucose metabolism using 2-deoxy-D-glucose (2-DG) or mitochondrial metabolism using oligomycin, allows determining functional metabolic profiles of cells based on their dependencies on these metabolic pathways.

Our analyses surprisingly showed no significant changes in overall substrate dependence—namely glucose dependence or FAO/AAO capacity—and dependence on ATP synthesis routes – namely mitochondrial or glycolytic - between chow *ad libitum* and HFD *ad libitum* conditions (Fig. 7c, 7d; 7g, 7h). This may partly be explained by the absence of fasting-induced stress in these conditions, as fasting may act as a stressor for neurons, leading to secondary effects on microglial metabolism [49]. However, during TRF, light-fed rats are fasted at ZT18, whereas dark-fed rats are fasted at ZT6, allowing assessment of microglial metabolic flexibility under fasting-induced stress. Here, CENCAT revealed differentially regulated microglial metabolism by TRF dependent on which diet was fed. In chow-fed rats, TRF primarily influenced substrate utilization, whereas in HFD-fed rats, TRF predominantly affected the route of ATP synthesis (Fig. 7e, 7f, 7i, 7j). Furthermore, dark phase TRF significantly increased glucose dependence and glycolytic capacity of microglia during the active phase (ZT18) in HFD-fed rats in comparison with chow-fed rats (Fig. 7i, 7j). These results suggest that metabolic flexibility is perturbed in microglia from HFD-fed rats, as microglia in chow-fed rats more readily switch to fatty acid and/or amino acid oxidation during feeding, while microglia in HFD-fed rats more heavily rely on non-mitochondrial glucose metabolism during feeding. However, linear regression analysis interestingly showed that the dependence on the metabolic route for ATP synthesis is correlated with body weight both at ZT6 and ZT18, and more strongly with fat mass only at ZT6 (Extended Data Fig.6), suggesting these TRF-induced mitochondrial metabolic alterations in HFD-fed rats may have beneficial secondary effects on body weight management. Interestingly, while the abundance of circulating classical and non-classical monocytes was affected, the impact of TRF on their metabolic profiles was markedly lower compared to microglia (Extended Data Fig. 5b for monocytes gating strategy, Extended Data Fig.7). This may further support the notion that the effects of TRF on microglial metabolism are secondary to its effects on neuronal activity, rather than a direct effect of feeding status. Taken together, we observed that TRF positively influences body composition and significantly affects the rhythmic expression of microglial genes. However, the detrimental effects of an HFD on microglial immunometabolism remain unaffected by TRF.

## Discussion

Our study aimed to elucidate the impact of time-restricted feeding on the microglial circadian clock and microglial metabolism during diet-induced obesity. We found that time-restricted feeding has beneficial effects on body composition and markedly impacted microglial gene rhythmicity. However, the detrimental effects of an HFD on microglial immunometabolism could not be reversed by time-restricted feeding.

In the presence of *ad libitum* access to food, HFD impairs circadian feeding rhythms more severely than does a standard chow diet [7, 50]. Therefore, rodents fed HFD *ad libitum* have a short fasting period and a long feeding window [5]. Such eating patterns disrupt circadian rhythms and the metabolic pathways that are coupled with the daily feeding rhythm. Emerging studies suggest that TRF may contribute to prevent metabolic dysfunctions and improve metabolic health [5, 51]. In the HFD condition, we found that TRF light- and dark-feeding caused similar alterations in body weight gain, weekly food intake, and weekly food efficiency. However, relative fat mass was only reduced in HFD dark-fed rats compared to HFD *ad libitum*, not in the HFD light-fed group[5]. This suggests that the timing of food intake during the active period, thus aligned with the sleep-wake rhythm, has a positive impact on body composition independent of calorie restriction *per se*. Our observation is supported by studies showing that HFD dark-fed mice had increased ghrelin and leptin effectiveness [52, 53], thereby contributing to less adiposity. Hence, the timing of food intake emerges as an important factor influencing body composition and metabolic dysfunctions.

Circadian rhythms and the immune system are intricately linked [25, 26]. Indeed, the majority of cells within the immune system exhibit autonomous circadian oscillations in gene expression, such as cytokines and receptors, in part under the control of the core clock machinery [26–31]. This has been shown to regulate immune cell trafficking, immune cell phenotypes, and inflammatory responses [28, 29, 31–34]. In addition to light exposure, also the timing of food intake influences circadian rhythms. For instance, timed feeding entrains rhythmic clock gene expression in the suprachiasmatic nucleus of mice kept in constant darkness and restores rhythmicity in mice kept in constant light [54, 55]. Moreover, in regular light/dark conditions, daytime feeding will shift the daily rhythm of many peripheral clocks [56, 57]. In the current study we found that microglial clock gene expression is highly sensitive to time-restricted feeding, while no significant effects were seen on mRNA levels of PBMCs core clock gene expression. A potential explanation for this discrepancy may be the blood–brain barrier, a morpho-functional structure that separates the CNS from the circulation, making the cellular microenvironment of microglia and PBMCs distinct [58]. Another possible explanation is the increased longevity and self-renewal capacity of microglia compared with PBMCs [59, 60]. The lifespan of human microglia is on average 4.2 years [59, 60], while PBMCs have a relatively short lifespan, typically lasting from a few days to a few weeks [61]. Therefore, the absence of clear TRF-induced alterations in PBMCs may be explained by their faster turnover and shorter lifespan, impeding sustained metabolic or epigenetic priming.

The basic paradigm of circadian regulation of metabolism is that rhythmic patterns of gene expression drive rhythms in cellular metabolism [5, 8]. TRF is known to increase the number of rhythmic transcripts in WT mice fed a chow diet [5]. In the chow condition, we observed that while the total number of genes gaining rhythmicity was similar between the light-fed and dark-fed rats, only ∼26% (326 of the 1,218) of genes that gained rhythmicity overlap between the chow light-fed group and dark-fed group. While most of the gene rhythms remained unchanged in the dark-fed group, in the chow light-fed group also a considerable number of genes lost rhythmicity. This indicates that the microglial intrinsic clock is highly sensitive to feeding cues [62]. Although HFD dark-feeding successfully rescued the loss of rhythmicity in some metabolism-related pathways, we did not see a reinstatement of gene expression in the electron transport chain genes. This indicates that (micro)environmental cues are more dominant in regulating microglial transcription compared to microglial intrinsic clock modulation.

Increased lipid content within cells is commonly associated with mitochondrial impairments via lipotoxicity, as has been well-documented in various cell types, including hepatocytes, muscle cells, and macrophages [63]. In hepatocytes, lipid overload can lead to impaired mitochondrial respiration and increased production of reactive oxygen species, contributing to conditions like non-alcoholic fatty liver disease [64]. Similarly, in muscle cells, lipid accumulation disrupts mitochondrial oxidative capacity, with implications for metabolic diseases like insulin resistance [65]. Macrophages exposed to lipid-rich environments also show mitochondrial dysfunction, compromising immune responses and promoting pro-inflammatory phenotypes [66]. Our findings may suggest a similar mechanism of lipid-induced mitochondrial and immunological dysfunction of microglia in HFD-fed rats, as genes related to lipid uptake and lipogenesis are upregulated, while microglia did not readily switch mitochondrial metabolism during feeding. However, these effects were not mitigated by TRF.

During inflammation, the microglial priming concept suggests that microglia exhibit heightened reactivity when faced with a similar second stimulus compared to those without prior stimulation [67–69]. This phenomenon implies a form of “microglial memory”, however, it is not fully understood whether this memory also involves immunometabolic reprogramming. While speculative, our data may indicate a potential “obesogenic memory” in microglia, as the metabolic impairment induced by HFD persists despite the absence of obesity in dark time-restricted feeding. This would imply two potential mechanisms: either microglial priming is facilitated by HFD-induced epigenetic alterations of microglia [70], or a microglial memory is stored elsewhere within the neural or glial network. For instance, it has been shown that a HFD-induced persistent elevation of microglial reactivity and consequent TNFα secretion induces mitochondrial stress in POMC neurons, which contributes to the development of obesity [40].

In summary, our study revealed that 4 weeks of time-restricted feeding during the active phase of the light/dark cycle effectively decreased fat gain induced by a high-fat diet. However, the HFD-induced impairment of cellular metabolism in microglia persisted. We propose an “obesogenic memory” in microglia, which we suggest as a potential explanation for weight cycling, *i.e.* the repeated loss and regain of weight often seen during dieting and known as the “yo-yo” effect [71, 72]. We think this new concept of ‘microglial obesogenic memory’ and the possibility that it hampers the effectiveness of long-term TRF and/or other caloric restriction strategies needs further investigation and prompts a re-evaluation of microglial functions in both health and disease contexts.

## Data availability

All data associated with this study are present in the paper or the Supplementary Materials. Any additional information required to reanalyse the data reported here is available from the corresponding authors upon reasonable request.

## Code availability

All codes used in the current study are available via GitHub (https://github.com/NeuroAnalyz/SpikeTrains-LFP) or from the corresponding authors upon reasonable request.

## Acknowledgments

We would like to thank Aldo Jongejan and Perry Moerland from the Bioinformatics Unit at the Amsterdam UMC for help and advice. We are also grateful to Richard Volckmann from the R2 Genomics Platform at the Amsterdam UMC for assistance and discussions.

## Material and methods

### Animals

Male Wistar rats (n = 293) (Charles River Laboratories, Sulzfeld, Germany) were group-housed in a 12h-12h light-dark (LD) or reversed 12h-12h dark-light (DL) cycle with lights on or off at 07:00 and 19:00 [with lights on defined as Zeitgeber Time zero (ZT0) and lights off as ZT12] and a room temperature of 22 ± 2°C. Upon arrival, animals received a standard chow diet (2018, Teklad diets, Invigo) or an obesogenic diet (D12331, Research Diets Inc). During an acclimatization period of 4 weeks, animals had *ad libitum* access to food and water. Animals were group-housed in different batches for logistic reasons. After the 4-week acclimatization period, animals receiving the same diet were randomized based on their body weight and the time-restricted feeding protocol was started. Animals in the LD room received light-phase feeding (10 hours of feeding from ZT1-ZT11) and animals in the DL room dark-phase feeding (10 hours of feeding from ZT13-ZT23). Animals fed *ad libitum* were randomized between the normal LD and reversed DL conditions. Body weight and food intake were monitored twice per week. Body composition of all animals was measured by MRI in the last week. All studies were approved by the Animal Ethics Committee of the Royal Dutch Academy of Arts and Sciences and performed according to the guidelines on animal experimentation of the Netherlands Institute for Neuroscience.

### Tissue sampling

After 4 weeks of TRF, animals were sacrificed at six different time points during the 24-hour day-night cycle (starting at ZT2, every four hours) by euthanasia with 60% CO_2_/40% O_2_ or 5% isoflurane, followed by decapitation. Trunk blood was collected, and the brain was isolated and dissected as described earlier [73]. One half of the forebrain was immediately stored in 4% PFA for 48 hours and transferred to 1x PBS containing 0.05% sodium azide (NaN_3_) for further analysis. From the other half of the forebrain, the hypothalamus and hippocampus were extracted, immediately frozen in liquid nitrogen and stored at −80°C for further analyses.

### Immunohistochemistry and Immunofluorescence

After 48 hours post-fixation the brains were equilibrated in a 30% sucrose solution. Coronal sections of the brains, with a thickness of 30_μm, were obtained using a cryostat (Leica Biosystems). These sections were collected and rinsed in 0.1_M Tris-buffered saline (TBS) to ensure proper tissue preparation.

Immunohistochemical staining was performed to detect the presence of Iba1 in the brain. Sections were incubated with a primary antibody, rabbit anti-Iba1 (1:1000, Synaptic Systems, No.234003, Germany). The primary antibodies were incubated overnight at 4_°C. Following rinsing, the sections were incubated with biotinylated secondary antibody, goat anti-rabbit IgG (1:400, BA-1000, Vector Laboratories, Inc., Burlingame, CA), and then further incubated in avidin-biotin complex (1:800, PK-4000, Vector Laboratories, Inc., Burlingame, CA) for 1_hour. The reaction product was visualized by incubating the sections in a solution of 1% diaminobenzidine with 0.01% hydrogen peroxide for 10_minutes. Subsequently, the sections were mounted on slides, dried, dehydrated through a graded series of ethanol, cleared in xylene, and coverslipped for image collection and quantification.

Images were acquired using a Zeiss AXIO Scope A1 microscope equipped with Plan-APOCHROMAT × 10/0.45 objective lenses and a Zeiss AxioCam MRc camera. AxioRel Vision 4.8 acquisition software was used for image capture. Quantification of imaging data was carried out in a blinded manner, where Iba1-immunoreactive (Iba1-ir) cell numbers and processes were manually counted. For immunofluorescence staining, fluorescent-conjugated antibodies Guinea pig anti-IBA1 (1:500, Synaptic System, No. 234 308) were added. After several rinses, brain sections were mounted on Superfrost Plus microscope slides, covered with cover slips, and stored at 4°C. Imaging was performed using a confocal microscope (TCS-SP8, Leica Biosystems, Heidelberg, Germany). The laser power, photomultiplier gain, and offset were maintained constant for all images.

### Blood mononuclear cell isolation and plasma collection

Trunk blood was used for PBMC isolation and plasma collection. For plasma collection, 2 ml blood was centrifuged for 15 min (4,000 rpm, 4°C, brake 9/9). Plasma was collected and stored at −20°C until further analyses. For PBMC isolation, 30 ml lysis buffer (containing 1 × ACK; 155 mM NH4Cl; 10 mM KHCO3; 0.1 mM EDTA) was added to ∼3 ml blood, mixed by gently vortexing and incubated at RT for 10–15 min. Cell suspension was centrifuged for 5 min (200 g, RT, brake 9/9). Supernatant was discarded and cells were resuspended in 2 ml PBS-FBS (PBS containing 1% FBS). After centrifugation of the new cell suspension for 5 min (200 g, RT, brake 9/9), supernatant was discarded, and cells were resuspended in 0.5 ml PBS-FBS. The obtained cell suspension was added to 4.5 ml RPMI medium and slowly layered on 5 ml Ficoll® (17-1440-02, GE Healthcare, Sigma-Aldrich®) in a 15 ml Falcon tube, followed by centrifugation for 30 min (400 g, 20°C, brake 1/1). The fuse interphase, containing PBMCs, was collected in 8 ml 1x HBSS, and centrifuged for 5 min (200 g, RT). Supernatant was discarded and the pellet was resuspended in RNA Lysis buffer (Qiagen) and stored at −80°C until RNA extraction. For CENCAT, PBMCs were isolated using LeucoSep tubes (Greiner). 8 ml blood was added to LeucoSep tubes containing 15 ml Ficoll, and PBS was added up to 50 ml. Tubes were centrifuged for 15 minutes (800 g, 20°C, brake 7/1). PBMCs were collected by pouring the supernatant into a new tube and washed twice with cold PBS after centrifugation (400 g, 4°C, brake 7/9). Cells were counted using a haemocytometer and up to 5×10^5^ cells per well were plated in a 96 well V-bottom plate for CENCAT.

### RNA isolation from isolated PBMCs and quantitative reverse-transcription polymerase chain reaction

Total RNA was isolated from PBMCs using a RNeasy Micro Kit (QIAGEN, Hilden, Germany, cat. no. 74004) following the manufacturer’s recommendations. We used 250 ng of RNA to generate cDNA with a Transcriptor First Strand cDNA Synthesis Kit (Roche, Basel, Switzerland, cat. no. 04897030001) following the manufacturer’s instructions. Quantitative PCR (qPCR) was performed using a SensiFAST™ SYBR® No-ROX Kit (Roche Bioline, cat. no. BIO-98020). Expression levels of all genes were normalized to the geometric mean of housekeeping genes. Primer sequences (see Table S1) were designed using the Basic Local Alignment Search Tool (BLAST) from the National Center for Biotechnology Information (NCBI). Primers were purchased from Sigma-Aldrich® and validated by melt curve analysis and DNA band size and/or purity on agarose gel electrophoresis (data not shown).

**Table S1.**
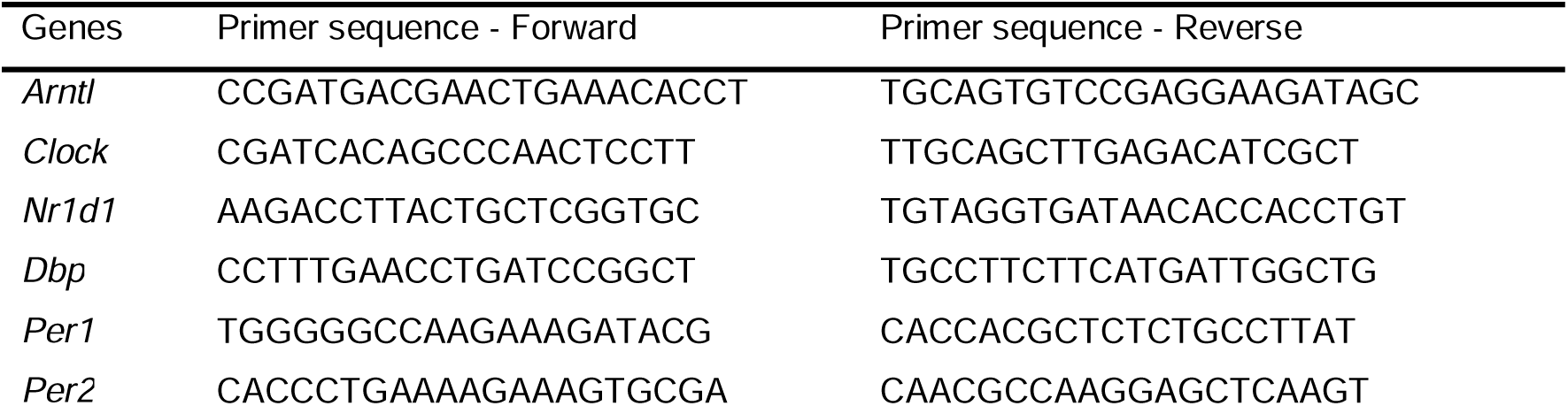
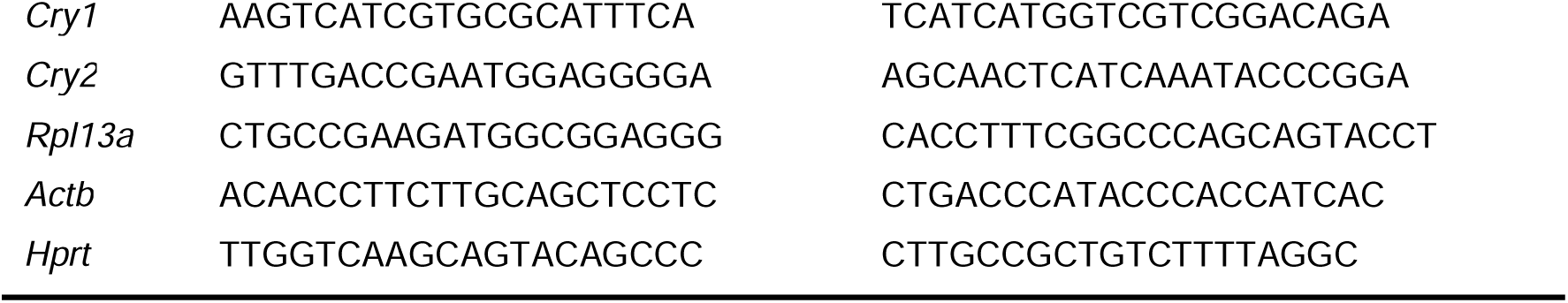
Primer sequences of target genes.

### Microglia isolation

Percoll isopycnic isolation was performed to isolate microglial cells from the cerebral brain cortex and one half of the midbrain without the hippocampus and hypothalamus. Briefly, brains were mechanically homogenized with RPMI 1640 medium (11875-093, Gibco™) using a tissue grinder (Mexico, Sigma-Aldrich), or digested in 0.2 mg/mL Collagenase IV (sigma, Cat# C5138) and 0.05 mg/mL DNAse I (Roche, Cat# 10104159001) in RPMI 1640 medium for 1 hour at 37°C, after which the brain homogenate was filtered through a 70 μm cell strainer (431751, Corning®) in a 50 ml Falcon tube. The filtered brain homogenate was centrifuged for 5 min (380 g, 4°C, brake 9/9). Supernatant was discarded and pellet was resuspended in 7 ml RPMI medium and mixed with 100% Percoll solution [for 10 ml: 9 ml Percoll^®^ stock (17-5445-01, GE Healthcare, Sigma-Aldrich®) together with 1 ml 10x HBSS (14185-045, Gibco™)]. The cell suspension was slowly layered on 70% Percoll solution [for 10 ml: 7 ml 100% Percoll solution with 3 ml 1x HBSS (14175-053, Gibco™)] in a 15 ml Falcon tube and centrifuged for 30 min (500 g, 18°C, brake 1/0). Myelin and cell debris were discarded and the fuse interphase, containing microglial cells, was carefully collected in 8 ml 1x HBSS, followed by centrifuging for 7 min (500 g, 18°C, brake 9/9). Supernatant was discarded and the microglial cell pellet was resuspended in RNA Lysis buffer (Qiagen) and stored at −80°C until RNA extraction. Alternatively, the microglial cell pellet was counted using a haemocytometer and up to 2.5×10^5^ cells per well were plated in a 96 well V-bottom plate for CENCAT.

### CENCAT

CENCAT was performed as described previously [48]. βES.HCl was synthesized as previously described [74] and dissolved in equimolar NaOH (200 mM) to neutralize the pH. Samples were split into four conditions in uniplo: control (DMSO+H_2_O vehicle control), 2-deoxy-D-glucose (2-DG, 100 mM), oligomycin (1 μM) and a combination of 2-DG and oligomycin. After plating in complete RPMI (cRPMI; RPMI1640 supplemented with GlutaMAX and 10% FCS), cells were incubated with metabolic inhibitors (10x work stock solutions in cRPMI) for 15 minutes at 37°C/5%CO_2_. All conditions were corrected for DMSO and H_2_O content. 500 μM βES was subsequently added (from a 10x work stock in cRPMI) and was allowed to incorporate into nascent proteins for 30 minutes at 37°C/5%CO_2_. After 30 minutes, cells were pelleted via centrifugation (500 g, 3 minutes), washed with PBS and incubated with Zombie Aqua (for microglia) or Zombie NIR (for PBMCs) fixable viability stain for 5 minutes at room temperature in the dark. After washing with PBS, microglia were incubated for 10 minutes at 4°C in the dark with a staining mix containing anti-RT1B-biotin, and fluorochrome-conjugated anti-CD86 and anti-CD32 antibodies in FACS buffer (1%BSA and 2 mM EDTA in PBS) supplemented with True Stain Monocyte Blocker solution (Biolegend) to prevent aspecific binding of fluorochrome-conjugated antibodies to myeloid cells. PBMCs were incubated with anti-RT1B-biotin and anti-CD32. Cells were incubated with these antibodies before fixation, as these were previously identified to generate a poor staining resolution after formaldehyde fixation (data not shown). Following washing with FACS buffer and centrifugation, cells were fixed using 2% formaldehyde in PBS for 15 minutes at room temperature in the dark. After washing with FACS buffer, cells were stored in FACS buffer wrapped in aluminum foil at 4°C, and fluorescent click labelling of βES and measurement was performed within 3 days.

For fluorescent click labelling, cells were first permeabilized using 1X permeabilization buffer (Invitrogen) for 15 minutes at room temperature in the dark. Next, cells were washed with click buffer (100 mM Tris-HCl, pH 7.4), and βES was labelled through a copper(I)-catalyzed azide-alkyne cycloaddition (CuAAC) reaction, using the following reaction mix in click buffer for 30 minutes at room temperature in the dark: 0.5 mM Cu(II)SO_4_, 10 mM sodium ascorbate, 2 mM THPTA and 0.5 μM AF405 Azide Plus. Subsequently, after centrifugation and washing with FACS buffer, microglia were incubated with a staining mix containing fluorochrome-conjugated streptavidin, and anti-CD200R, anti-CD45 and anti-CD11b/c antibodies in FACS buffer supplemented with True Stain Monocyte Blocker solution. PBMCs were incubated with fluorochrome-conjugated streptavidin, and anti-CD3, anti-His48, anti-CD43, anti-CD45 and anti-CD161 antibodies. After 15 minutes incubation at room temperature in the dark, cells were centrifuged, resuspended in FACS buffer, and acquired on a CytoFLEX S cytometer (Beckman Coulter).

Rats were sacrificed in two block of three consecutive days. Quality control within each block, by comparing the effects of inhibitors between days, revealed one day of technical variation where inhibitory effects of oligomycin (for microglia) or 2-DG (for PBMCs) was significantly less compared to the other sacrifice days. Thus, all data from those days was omitted, resulting in a sample size of 4-6 rats per group.

Antibody information is provided in Table S2.

| Reagent/antibody | Source | Identifier |
| --- | --- | --- |
| Anti-rat CD11b/c-PE-Cy7 | Biolegend | 201817 |
| Anti-rat CD161-APC | Biolegend | 205606 |
| Anti-rat CD200R-FITC | Biolegend | 204905 |
| Anti-rat CD32 | BD Biosciences | 550270 |
| Anti-rat CD32-APC-Vio770 | Miltenyi Biotec | 130-103-391 |
| Anti-rat CD3-VioGreen | Miltenyi Biotec | 130-119-125 |
| Anti-rat CD43-PE | Biolegend | 202812 |
| Anti-rat CD45-PE-Dazzle594 | Biolegend | 202223 |
| Anti-rat CD86-APC | Miltenyi Biotec | 130-109-130 |
| Anti-rat His48-FITC | eBiosciences | 11-0570-82 |
| Anti-rat RT1B-biotin | Miltenyi Biotec | 130-108-704 |
| Streptavidin-BV785 | Biolegend | 405249 |
| Zombie Aqua fixable viability stain | Biolegend | 423101 |
| Zombie NIR fixable viability stain | Biolegend | 423105 |

### Microglial RNA sequencing

To study differences in microglial gene expression, RNA sequencing was performed. RNA from microglial cells was isolated using the RNeasy Micro Kit (Qiagen), according to the manufacturer’s guidelines. mRNA was enriched from total RNA with the Dynabeads® mRNA Purification Kit (61006, Invitrogen). A strand-specific transcriptome library was constructed using the MGIEasy RNA Directional Library Prep Set (16 RXN) (1000006385, MGI). Briefly, after cDNA synthesis, adaptor ligation was performed to ligate adaptors with the cDNAs. Subsequently, a PCR reaction was conducted to amplify the cDNAs, after which the library quality was checked using the Agilent 2100 Bioanalyzer (G2939AA, Agilent). Finally, single-stranded cyclized products were produced from single-stranded PCR products. To perform sequencing, single-stranded circle DNA molecules were replicated via rolling cycle amplification and a DNA nanoball (DNB) was generated. Sequencing was performed through combinatorial Probe-Anchor Synthesis (cPAS) on the DNBSEQ-G400 (G400, MGI).

### Statistical analyses

Data are expressed as mean ± SEM. Two-way ANOVA with multiple comparisons was used to compare multiple groups at each weekly time point throughout the TRF period and at each ZT throughout the 24-hour day-night cycle. P < 0.05 was considered to be statistically significant. Statistical analyses were performed using GraphPad Prism (v 9.5.1).

### RNA-seq data analyses

Sequencing data was filtered using SOAPnuke [75] by (1) Removing reads containing sequencing adapter; (2) Removing reads whose low-quality base ratio (base quality less than or equal to 15) is more than 20%; (3) Removing reads whose unknown base (’N’ base) ratio is more than 5%, afterwards clean reads were obtained and stored in FASTQ format. Clean sequence reads were aligned to the reference genome (Rattus_norvegicus, GCF_000001895.5_Rnor_6.0) using HISAT2 70, after which Ericscript (v0.5.5) [76] and rMATS (V3.2.5) [77] were used to detect fusion genes and differential splicing genes (DSGs), respectively. Clean sequence reads were aligned to a reference gene set, a database built by the Beijing Genomic Institute in Shenzhen (BGI), in which known and novel coding and non-coding transcripts were included, using Bowtie2 [78]. Raw counts matrix was normalized by transcripts per million (TPM). We performed rhythm analysis using cosinorPy(v2.1) [79], MetaCycle [80] and CompareRhythm(v1.0.1) [44]. We described the circadian rhythmicity of genes within each group using 1 component trigonometric regression model using cosinor.fit_group in CosinorPy. The mean expression of each group was calculated for circadian comparison among groups and genes with mean expression less than 5.5 in each group were filtered. We applied model selection methods in compareRhythm using the Bayesian information criterion. Comparison of the amplitude (one-half the peak-to-trough difference) and acrophase (the peak time of the fitted curve) of selected genes among the groups was performed using cosinor1.test_cosinor_pairs. We applied GSVA (v1.48.3) [81] to estimate the variation of pathway activity within each group using the Poisson kernel. The canonical pathways gene sets were downloaded from the Molecular Signatures Database (MSigDB, https://www.gsea-msigdb.org/gsea/msigdb). Pathways were clustered using Mfuzz (v2.60.0) [82] according to their expression change over time and Metascape [http://metascape.org] web-based tool. CompareRhythms was also performed to detect the rhythm of pathways. Differential expression analysis was performed by limma (v3.56.2) [83]. We began by applying the *lmFit ()* function to fit a linear model. Subsequently, *contrast.fit ()* was utilized to compute estimated coefficients and standard errors, followed by *eBayes ()* to calculate moderated t-statistics, moderated F-statistics, and log-odds. Protein-protein interactions were predicted using STRING (https://string-db.org/) [84]. The chord plot was generated with the circlize package (v0.4.15), while the pathways network plot was created by igraph (v1.5.1). The Sankey plot was generated by ggsankey (v 0.0.99999). PREMANONA was used to find transcriptomic differences between groups by pairwiseAdonis(v 0.4.1) [85].

## Supplementary figure legends

**Extended Data Fig.1.**
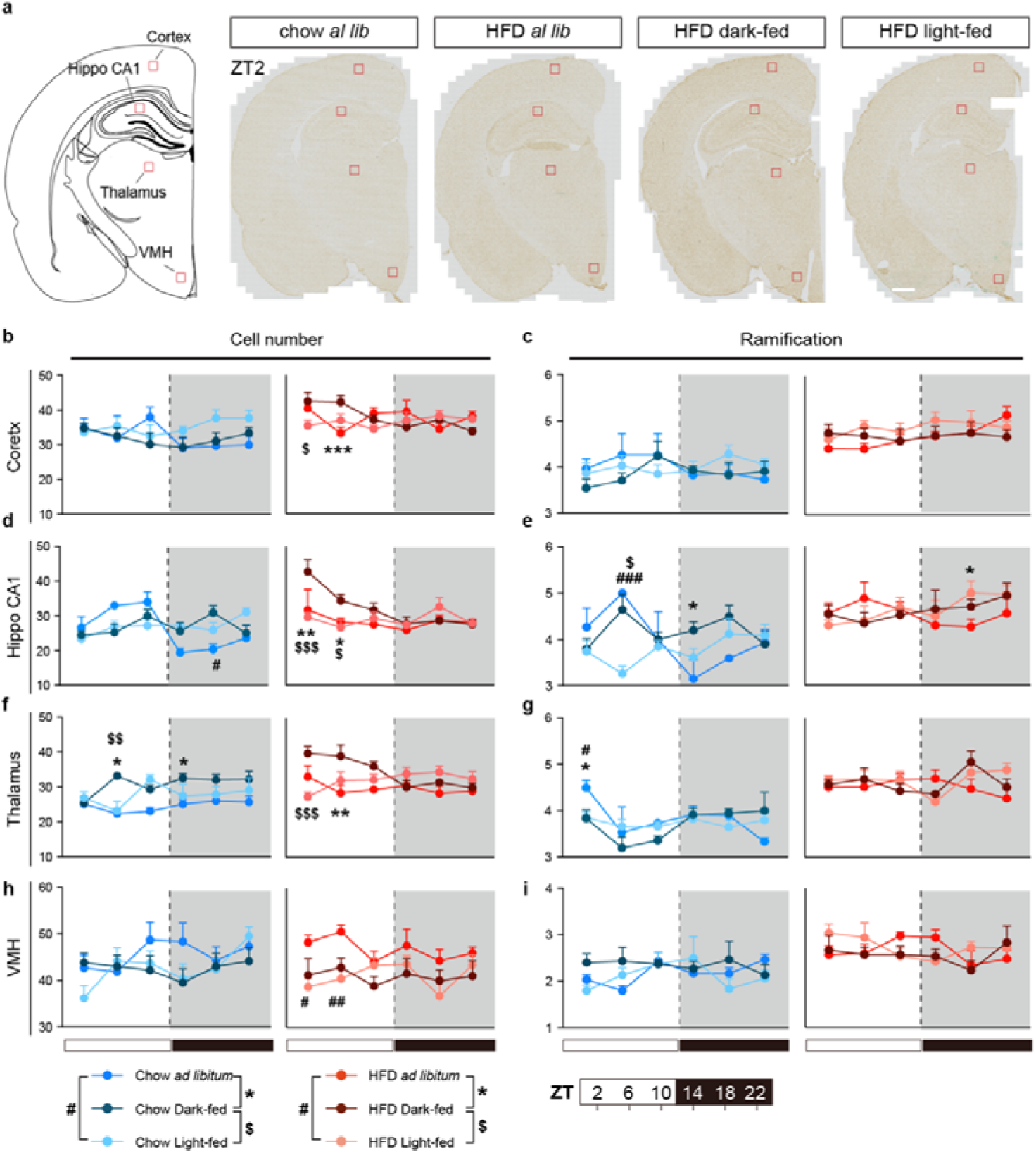
Microglial morphological assessment in different brain regions. a, Microglial morphology was determined in brain regions indicated with small red rectangles in the transversal brain section in different groups. **b-i,** Quantifications of microglial cell number and ramifications in the cortex (**b,c**), hippocamus CA1 (**d,e**), thalamus (**f,g**) and the ventromedial nucleus of the hypothalamus (VMH) (**h,i**). Data are shown as mean ± SEM and were analysed by two-way ANOVA with repeated measures (*, ^#, $^ p< 0.05, ^**, ##, $$^p<0.01, ^***, ###, $$$^p<0.001).

**Extended Data Fig.2.**
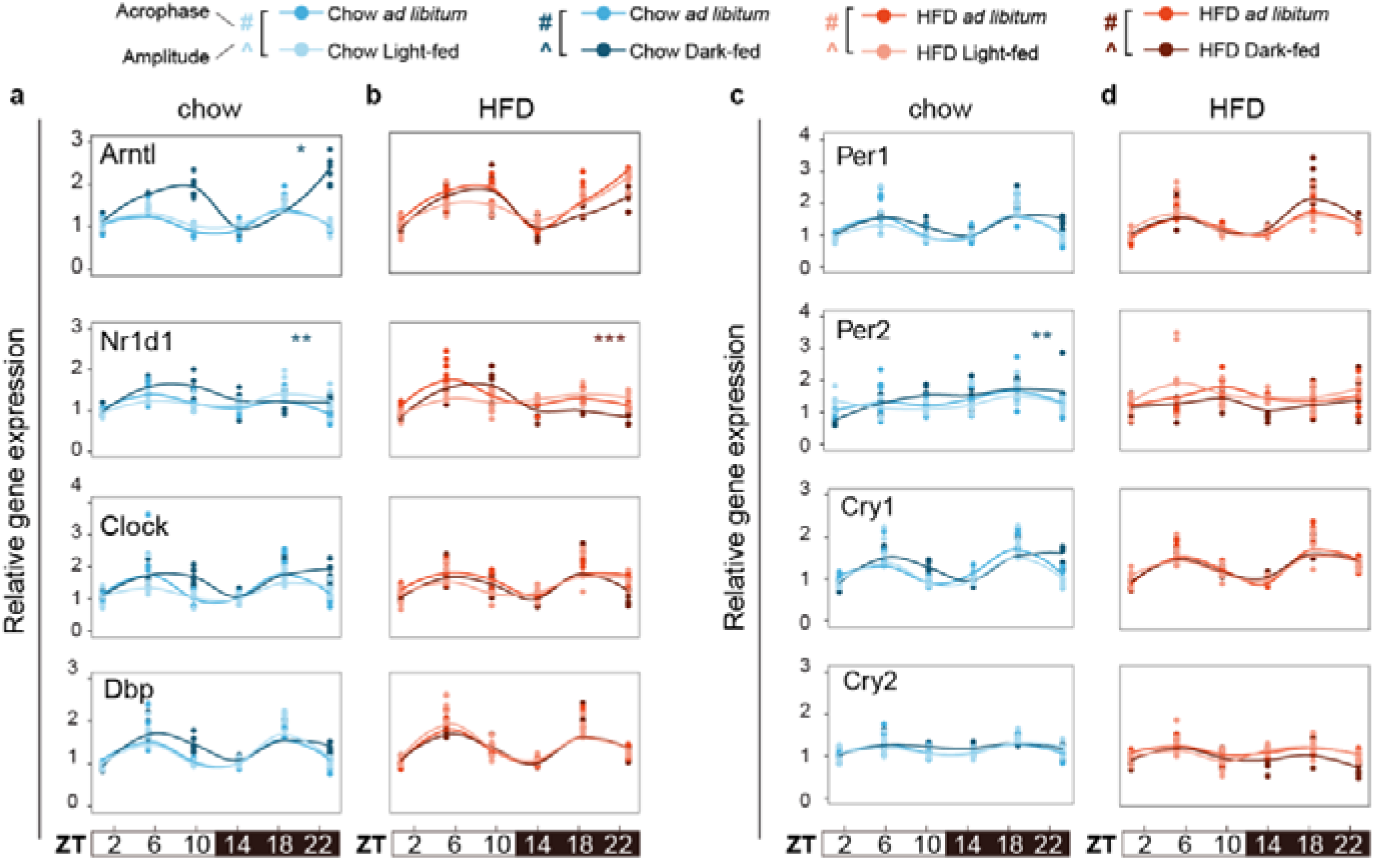
TRF effects on molecular clock gene expression in PBMCs a-d, Relative daily gene expression of 8 core clock genes during various feeding patterns under chow **(a,c)** and HFD **(b,d)** conditions (timepoints from ZT2 to ZT22, at 4-hour intervals). Coloured asterisks in the upper right corner indicate adjusted p-values [adj P] obtained from JTK_CYCLE analysis for different feeding patterns: *P < 0.05, **P < 0.01, ***P < 0.001.

**Extended Data Fig.3.**
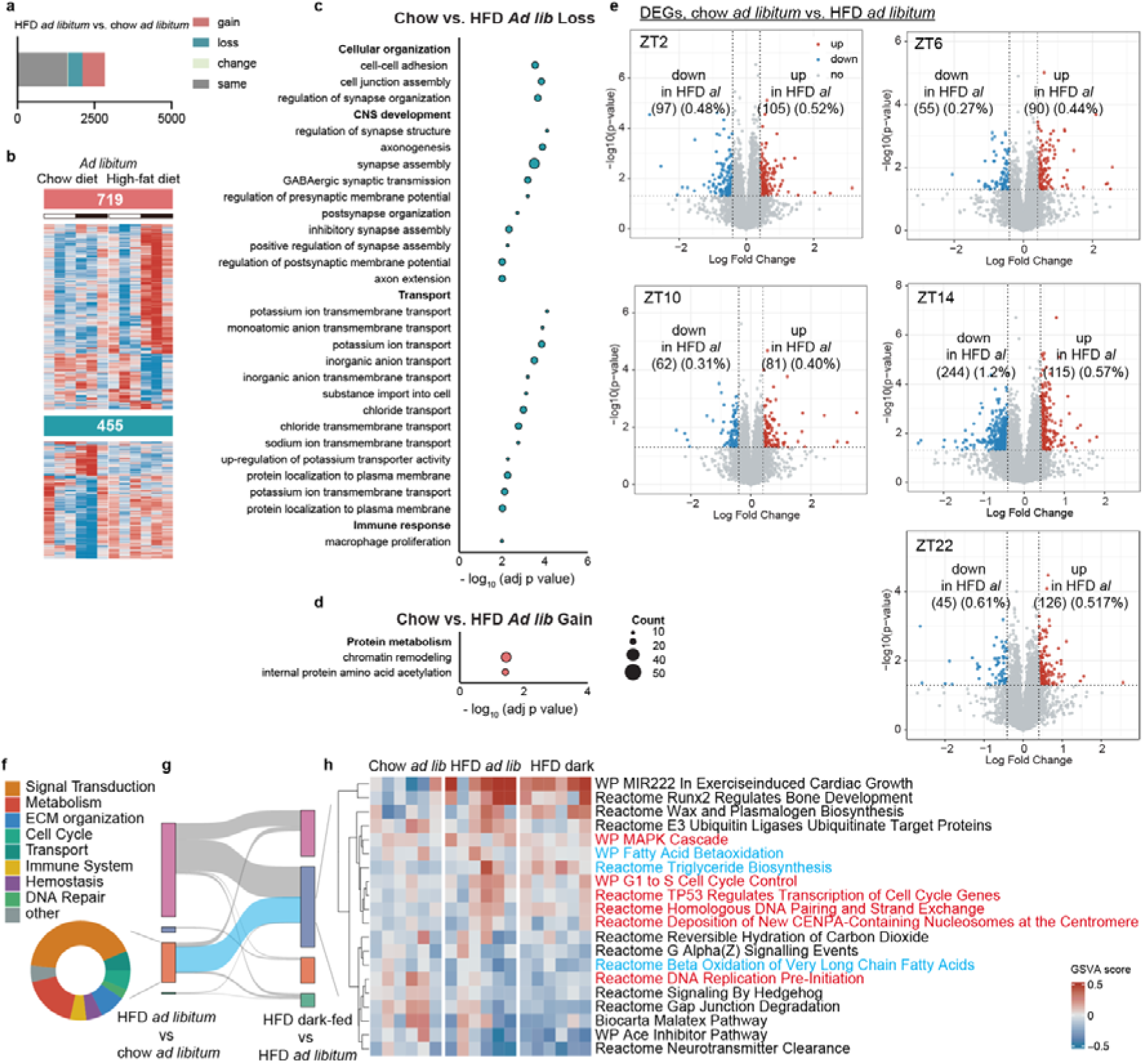
Enriched pathways in loss, gain categories based on gene ontology in chow vs. HFD *ad libitum*. a, (Stacked) bar graph depict the percentages of gained, lost, shifted, and unchanged genes relative to the total number of rhythmic genes in chow *ad libitum* vs HFD *ad libitum*. **b,** Heatmaps displaying the z-scored normalized transcript expression levels in chow *ad libitum* vs HFD *ad libitum*. **c-d,** Pathways involved lost (**c**) and gained (**d**) genes in the chow vs. HFD *ad libitum* comparison. Enlarged and bold terms serve as summaries of sub pathways. Dot size corresponds to the number of genes associated with each pathway, and log_10_ adjusted p-value thresholds are shown on the x-axis. **e,** Volcano plots showing differentially expressed genes (DEGs) at different time points under chow *ad libitum* and HFD *ad libitum* conditions. Genes with significant changes (pD<D0.05) are indicated in red (fold change ≥D2) or blue (fold changeD≤D-2). **f,** Pathway categories involved in genes with gained rhythmicity in HFD *ad libitum* vs. chow *ad libitum.* **g,** Sankey diagram illustrates alternations in gain, loss, shift, and unchanged categories between different comparison groups. **h,** Enriched pathways, based on the WikiPathways (WP), Reactome Pathway and Biocarta Pathways databases, depicted are pathways related to the transition from gained rhythmicity in HFD vs. chow *ad libitum* to lost rhythmicity in HFD dark-fed group vs. *ad libitum* group. Pathways highlighted in red are specifically associated with the cell cycle.

**Extended Data Fig.4.**
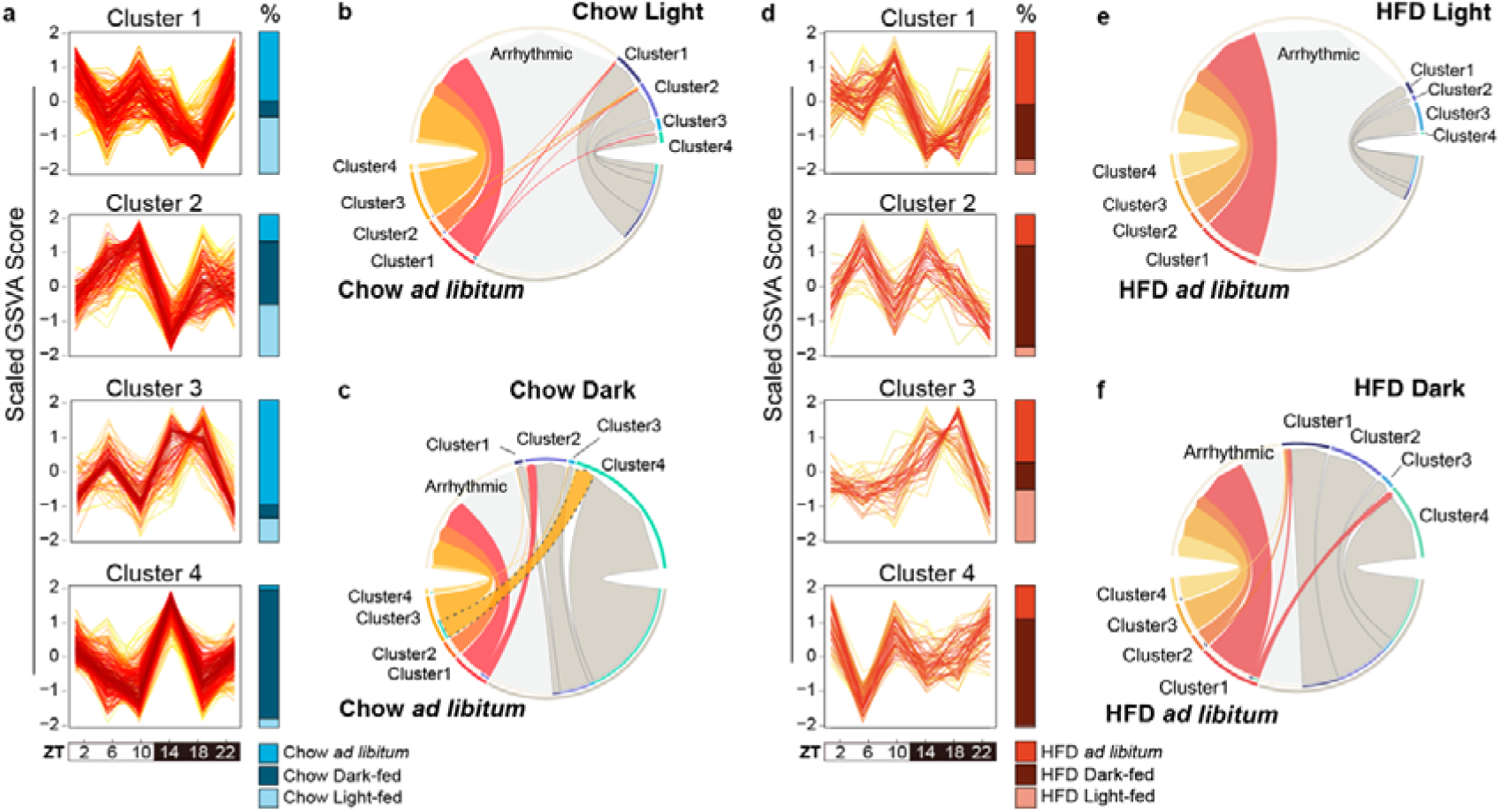
HFD reduces the number of clustered pathways. a, The Mfuzz algorithm was applied to cluster pathways resulting in four distinct clusters. The blue bars on the right represent the proportion (%) of signaling pathways within the same cluster across different TRF groups in chow. **b,c,** Chord plot illustrates the alternations of the 4 clusters of pathways between chow light-fed vs. *ad libitum*-fed (**b**), and chow dark-fed vs. *ad libitum*-fed (**c**). The colours at the edge of the circles indicate different clusters. **d,** The Mfuzz algorithm illustrates four pathway clusters across all feeding patterns in HFD-fed rats. The red bars on the right represent the proportion (%) of signaling pathways within the same cluster across different TRF groups in HFD. **e,f,** Chord plot illustrates the alternations of the 4 clusters between HFD light-fed vs. *ad libitum* (**e**), and HFD dark-fed vs. *ad libitum* (**f**). The colours at the edge of the circles indicate different clusters.

**Extended Data Fig.5.**
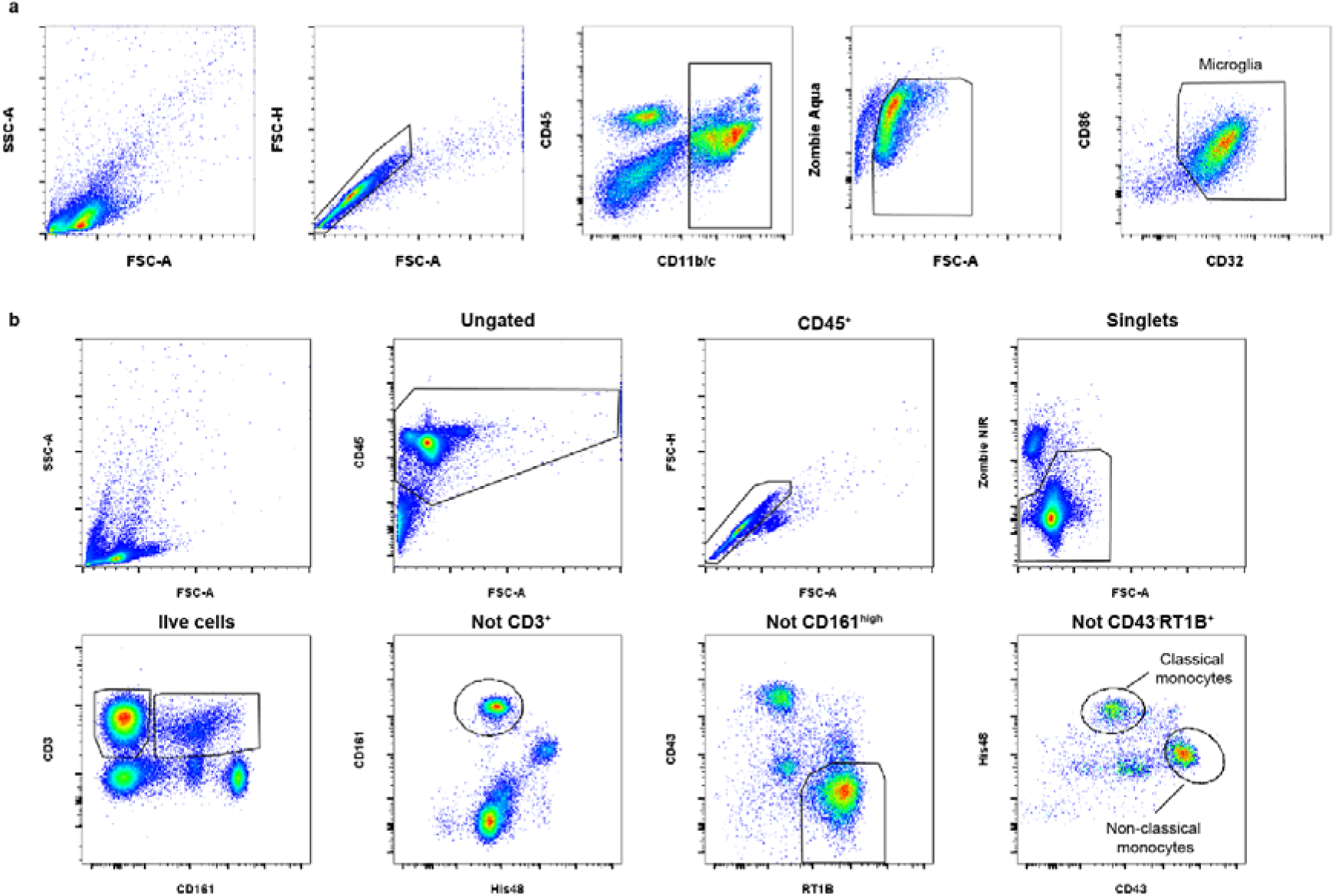
Flow cytometry gating strategies. **a**, Gating strategy for detecting microglia by flow cytometry. After excluding doublets and selecting CD11b/c^+^ cells, dead cells were excluded using the viability stain Zombie Aqua, and microglia selected based on co-expression of CD86 and CD32. **b**, Gating strategy for detecting circulating monocyte subsets by flow cytometry. After selecting CD45^+^ leukocytes and excluding doublets and dead cells, classical monocytes were defined as CD3^-^CD161^-^RT1B^low^CD43^low^His48^+^. Non-classical monocytes were defined as CD3^-^CD161^-^RT1B^low^CD43^++^His48^-^. Antibody information is provided in Table S2.

**Extended Data Fig.6.**
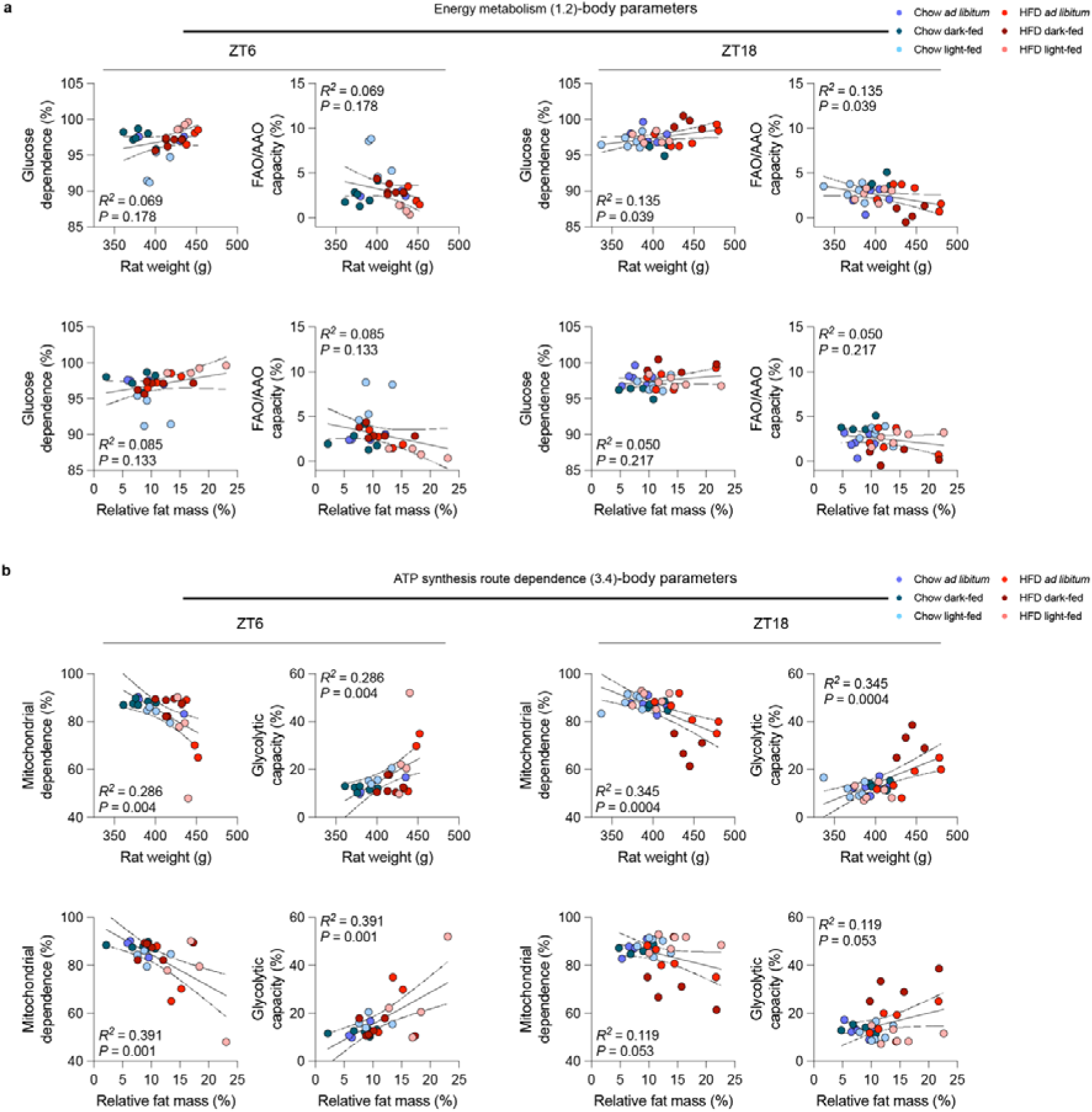
Linear regression analysis of CENCAT parameters with metabolic parameters a, Regression analysis between substrate dependence and metabolic parameters in all groups at ZT6 and ZT18. **b,** Regression analysis between ATP synthesis route dependence and metabolic parameters in all groups at ZT6 and ZT18.

**Extended Data Fig.7.**
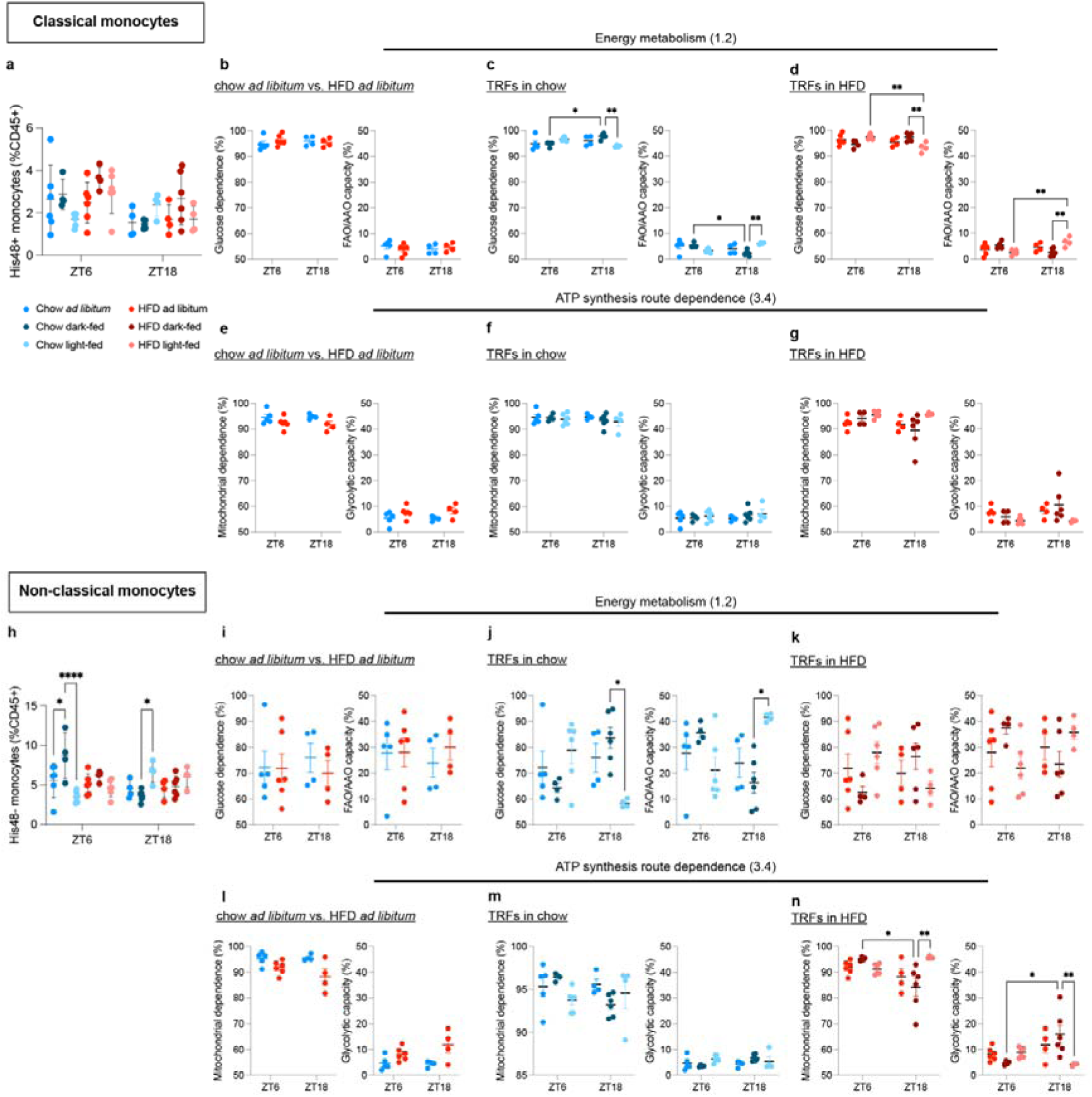
Metabolic profiling of circulating classical and non-classical monocytes by CENCAT a, Proportion of classical and non-classical monocytes in PBMCs. **b-d**, Energy metabolism of microglia using CENCAT from chow *ad libitum* vs. HFD *ad libitum*-fed rats (**b**), different TRFs in chow diet-fed rats (**c**) and different TRFs in HFD-fed rats (**d**) at ZT6 and ZT18 in classical monocytes. **e-g**, Dependence on route of ATP synthesis of microglia from chow *ad libitum* vs. HFD *ad libitum-*fed rats (**f**), different TRFs in chow diet-fed rats (**g**) and different TRFs in HFD-fed rats (**h**) at ZT6 and ZT18 in classical monocytes. **h,** Proportion of non-classical monocytes in PBMCs. **i-k**, Energy metabolism of microglia using CENCAT from chow *ad libitum* vs. HFD *ad libitum*-fed rats (**i**), different TRFs in chow diet-fed rats (**j**) and different TRFs in HFD-fed rats (**k**) at ZT6 and ZT18 in classical monocytes. **l-n**, Dependence on route of ATP synthesis of microglia from chow *ad libitum* vs. HFD *ad libitum-*fed rats (**l**), different TRFs in chow diet-fed rats (**m**) and different TRFs in HFD-fed rats (**n**) at ZT6 and ZT18 in classical monocytes.

